# Phase-locked dual-band vibration bursts strengthen anti-kindling by vibrotactile coordinated reset stimulation in a spiking network model

**DOI:** 10.64898/2026.11.05.764557

**Authors:** Shreya Abhijit Patki, SKM Varadhan

## Abstract

Parkinson’s disease and related movement disorders are accompanied by pathologically synchronized neuronal activity in cortico-basal ganglia circuits. Coordinated reset (CR) stimulation was derived computationally as a means of suppressing such activity at its source, by resetting the phases of the subpopulations of a synchronized network at staggered times. Under spike-timing-dependent plasticity (STDP), the resulting decorrelated firing depresses excitatory coupling and establishes a weakly coupled, desynchronized state that outlasts stimulation, a process termed anti-kindling. Sustained post-stimulation benefit has been demonstrated for electrical CR stimulation in parkinsonian primates and in proof-of-concept patient studies, and vibrotactile CR (vCR) stimulation realizes the same principle non-invasively through bursts of fingertip vibration. The deployed vCR protocol employs a carrier frequency of 250 Hz, which preferentially recruits Pacinian afferents. Because Pacinian receptive fields extend substantially beyond the stimulated digit, inputs originating at adjacent fingertips converge on overlapping regions of the target population, which degrades the spatial separation on which CR depends. Here we examine whether the spectral composition of the individual burst, independent of the stimulation schedule, governs the magnitude of anti-kindling. In a Hodgkin-Huxley network with STDP, driven through a two-channel model of fast-adapting afferents, a dual-band burst superimposing a 30 Hz component on the 250 Hz carrier yielded the lowest post-stimulation coupling among eleven vCR patterns, at one third of the carrier’s drive power. Recruitment of focal Meissner afferents by the 30 Hz component restricts part of the drive to the stimulated subpopulation, and phase locking of that component to the burst envelope generates post-burst pauses that reset the phase of the stimulated neurons. Spectral composition therefore determines both the spatial selectivity of vCR stimulation and its capacity for phase resetting.

**Author summary:** Excessive neuronal synchronization contributes to the symptoms of several brain disorders. Coordinated Reset (CR) stimulation was designed to counteract it by recruiting the capacity of neural circuits to reorganize their own connectivity. Stimuli delivered out of phase at several sites drive the affected neurons apart in time, and the synaptic changes that follow hold them apart once stimulation has ceased. Effects that persist in this way are what make a non-invasive form of the therapy realistic, since a device worn for a few hours can then carry benefit into the remainder of the day. Vibrotactile CR stimulation is a non-invasive form, and it reaches the nervous system through the mechanoreceptors of the fingertip. Computational work on it has so far treated the vibration burst as fixed and optimized the schedule on which bursts are delivered. We find that burst composition determines the magnitude of the lasting effect, that two separable mechanisms are responsible, one resting on the timing of the waveform and one on the class of skin receptor recruited. This can, in the future, be extended to pre-clinical and clinical studies.

## Introduction

Parkinson’s disease is the second most common neurodegenerative disorder. Its motor signs are accompanied by excessive synchronization of neuronal activity in the beta band (13-30 Hz) in the subthalamic nucleus and the wider cortico-basal ganglia-thalamocortical circuit [1,2]. This synchrony is linked to the severity of motor impairment. High-frequency deep brain stimulation of the subthalamic nucleus improves motor function in advanced disease [3] while simultaneously suppressing beta activity [4]. Deep brain stimulation is nonetheless invasive, and its benefit lapses once stimulation is withdrawn.

Coordinated reset (CR) stimulation was developed to counteract the synchrony itself instead of overriding it: brief stimuli delivered to several sites of a synchronized population in a staggered order reset the phases of its subpopulations at different times, so that they fire out of step with one another [5]. Because synaptic strength in these circuits depends on the relative timing of pre- and postsynaptic spikes [6], desynchronized firing also weakens the coupling that sustains synchrony. In networks with spike-timing-dependent plasticity (STDP), CR moves the network from a synchronized, strongly coupled state to a desynchronized, weakly coupled one in which it remains after stimulation ends; this lasting reduction of coupling is termed anti-kindling [7]. Anti-kindling has been demonstrated in networks of bursting subthalamic neurons [8], in a large-scale model of the subthalamic nucleus and globus pallidus [9] and in a ring network of Hodgkin-Huxley neurons with STDP, in which the CR rate and the stimulation intensity determine the strength of the effect [10]. In networks with spatially structured connectivity, the shape and amplitude of the individual stimuli also determine how strongly CR weakens synapses, and pronounced desynchronization during stimulation does not by itself guarantee a lasting effect [11]. It has been shown experimentally that the benefit of CR outlasts the stimulation itself. In parkinsonian primates, CR stimulation of the subthalamic nucleus produced motor improvement that persisted for days to weeks after stimulation ceased, which conventional high-frequency stimulation did not [12,13], and in a proof-of-concept study in patients it reduced beta activity cumulatively and improved motor function [14].

Vibrotactile CR (vCR) delivers the staggered stimuli non-invasively as bursts of vibration applied to the fingertips, extending the principle of CR from electrical to sensory stimulation [15–17]. The deployed protocol stimulates four fingertips of each hand with 100 ms bursts at a carrier frequency of 250 Hz, delivered at a CR rate of 1.5 Hz with random jitter of burst onsets and a cycle of three CR periods on and two off [18,19]. Open-label studies of this protocol have reported motor improvement that accumulated over months of use [19,20]. Computational work on CR has concentrated on the stimulation schedule, optimizing the CR rate and intensity, the order in which sites are addressed and the jitter of burst onsets [10,21,22], whereas the content of the individual stimulus has received less attention. Throughout this paper, "CR rate" refers to the 1.5 Hz repetition of the stimulation cycle and "carrier frequency" to the vibration frequency within a burst.

The glabrous skin of the hand is innervated by four classes of cutaneous mechanoreceptive afferent, each specialized for a distinct perceptual function [23,24]. Two of them respond at the frequencies used in vibrotactile CR, and they differ in both tuning and spatial reach. Fast-adapting type I afferents (FA1, Meissner corpuscles) respond best near 30 Hz and have small receptive fields whose sensitivity falls steeply outside a sharply bounded central zone [25–27]. Fast-adapting type II afferents (FA2, Pacinian corpuscles) respond best near 250 Hz and have a single zone of maximal sensitivity beyond which the threshold rises only gradually, so that one afferent still responds to indentation applied well away from its centre [26,28]. A 250 Hz carrier therefore recruits predominantly an afferent class whose fields extend beyond the stimulated fingertip. CR relies on stimulating separate subpopulations, and its efficacy depends on how far each stimulus spreads relative to the spacing of the stimulation sites [11,29]. Inputs that reach the target through large receptive fields should therefore overlap between neighbouring fingertips, and recruiting focal FA1 afferents should restore the separation between the stimulated subpopulations. Compound vibrotactile stimuli that combine FA1-targeting 32 Hz bursts at the fingertips with FA2-targeting 250 Hz bursts at the middle phalanx have been proposed on these physiological grounds [15], but their effect on anti-kindling has not been examined. The content of a burst may also act at the level of single neurons. CR desynchronizes by resetting the phase of each stimulated subpopulation [5], whereas a model of vibrotactile CR with leaky integrate-and-fire neurons was built on the premise that vibratory stimuli do not reset the phase of the neuronal oscillation [19]. Whether a burst resets phase is likely to depend on how the target neurons respond to transient input. Fast-spiking interneurons in somatosensory cortex have type 2 excitability: they cannot fire periodically below a critical frequency and, close to threshold, alternate between periodic firing and subthreshold oscillation [30]. The Hodgkin-Huxley neuron shares this behaviour, because repetitive firing begins at a Hopf bifurcation [31]. In such neurons, a burst whose drive returns smoothly to zero could leave the membrane close to its resting state, from which firing resumes only after a delay. A pause of this kind, locked to the end of each burst, would act as a phase reset of the stimulated subpopulation. Because FA1 afferents fire in phase with each cycle of low-frequency vibration [25,32], the drive that a 30 Hz component delivers to the target follows the timing of its cycles, so that the alignment of these cycles with the burst envelope determines how the drive ends.

Here we test these two routes by comparing the clinical 250 Hz carrier with a dual-band burst in which a 30 Hz component and a 250 Hz component share a Gaussian envelope (Fig 1), together with nine further vibrotactile CR patterns and control conditions that separate dose, envelope, spatial selectivity and the phase of the 30 Hz component. The target is a ring network of Hodgkin-Huxley neurons with STDP [10] that receives the five-fingertip input through a two-channel model of FA1 and FA2 afferents (Fig 2). It represents a somatotopically organized population that has reached a synchronized, strongly coupled state; neighbouring neurons represent neighbouring skin locations, and its intrinsic rhythm near 70 Hz is a property of the model neurons rather than a representation of beta activity. The primary outcome is the lasting reduction of synaptic coupling after stimulation, with synchrony and the coherence between stimulated network sectors as secondary outcomes.

**Fig 1.**
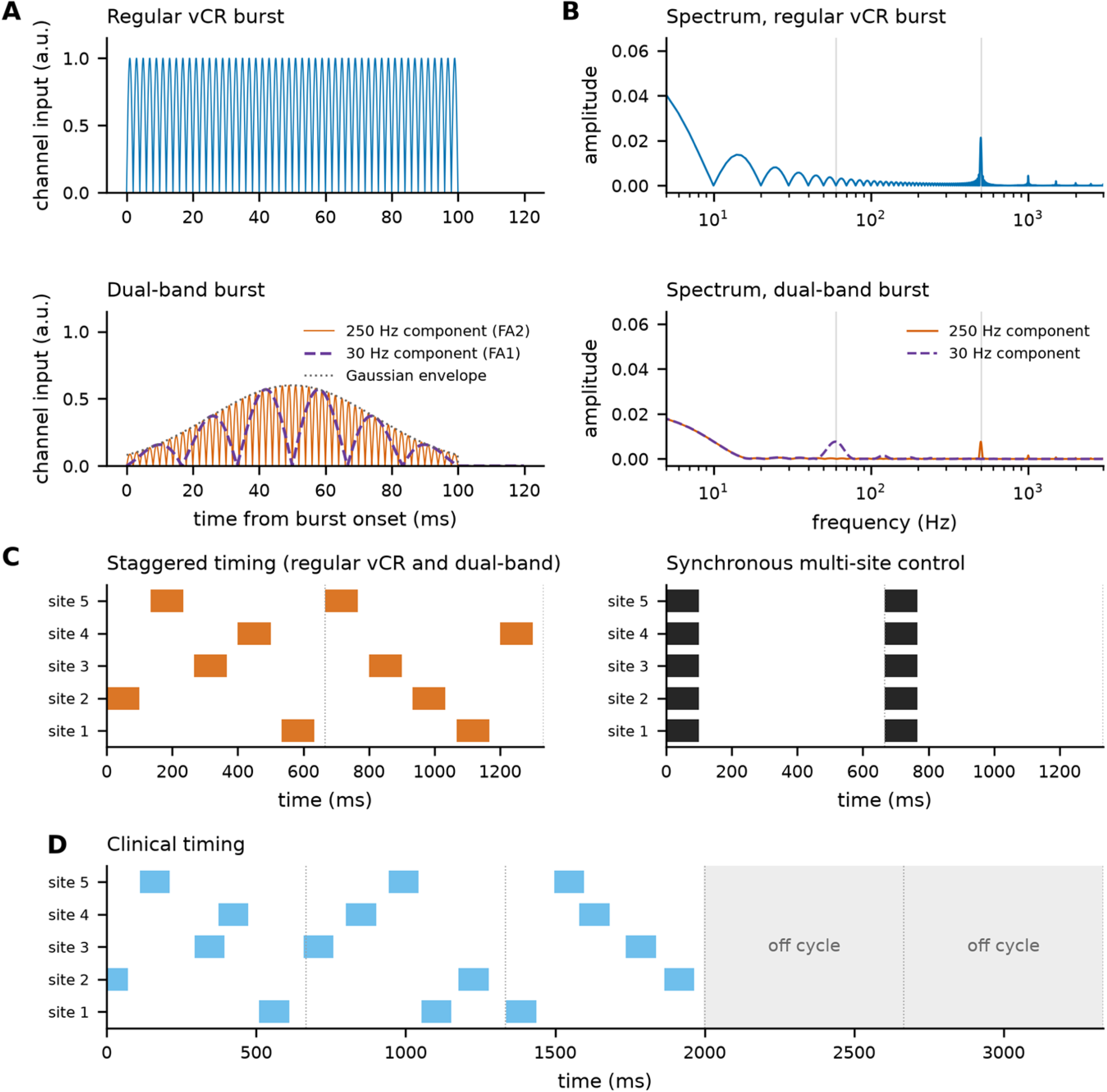
The dual-band burst keeps the CR timing of vibrotactile CR and changes the content of every burst. (A) Input delivered to each afferent channel by a single burst of regular vCR (250 Hz, rectangular envelope, amplitude 1) and by the dual-band burst (30 Hz component to FA1, dashed; 250 Hz component to FA2, solid; Gaussian envelope with SD 25 ms, dotted; amplitude 0.6 each). The 30 Hz component completes three cycles and reaches zero at burst offset together with the envelope. (B) Amplitude spectra of the bursts in (A). Full-wave rectification shifts each component to twice its frequency (grey lines at 60 and 500 Hz) and adds a low-frequency component set by the envelope. (C) Site timing over two 667 ms CR cycles (dotted lines) for staggered stimulation, shared by regular vCR and the dual-band burst, and for the synchronous multi-site control. (D) Timing of the clinical-style protocol over five cycles, with random onset jitter and three cycles on, two off.

**Fig 2.**
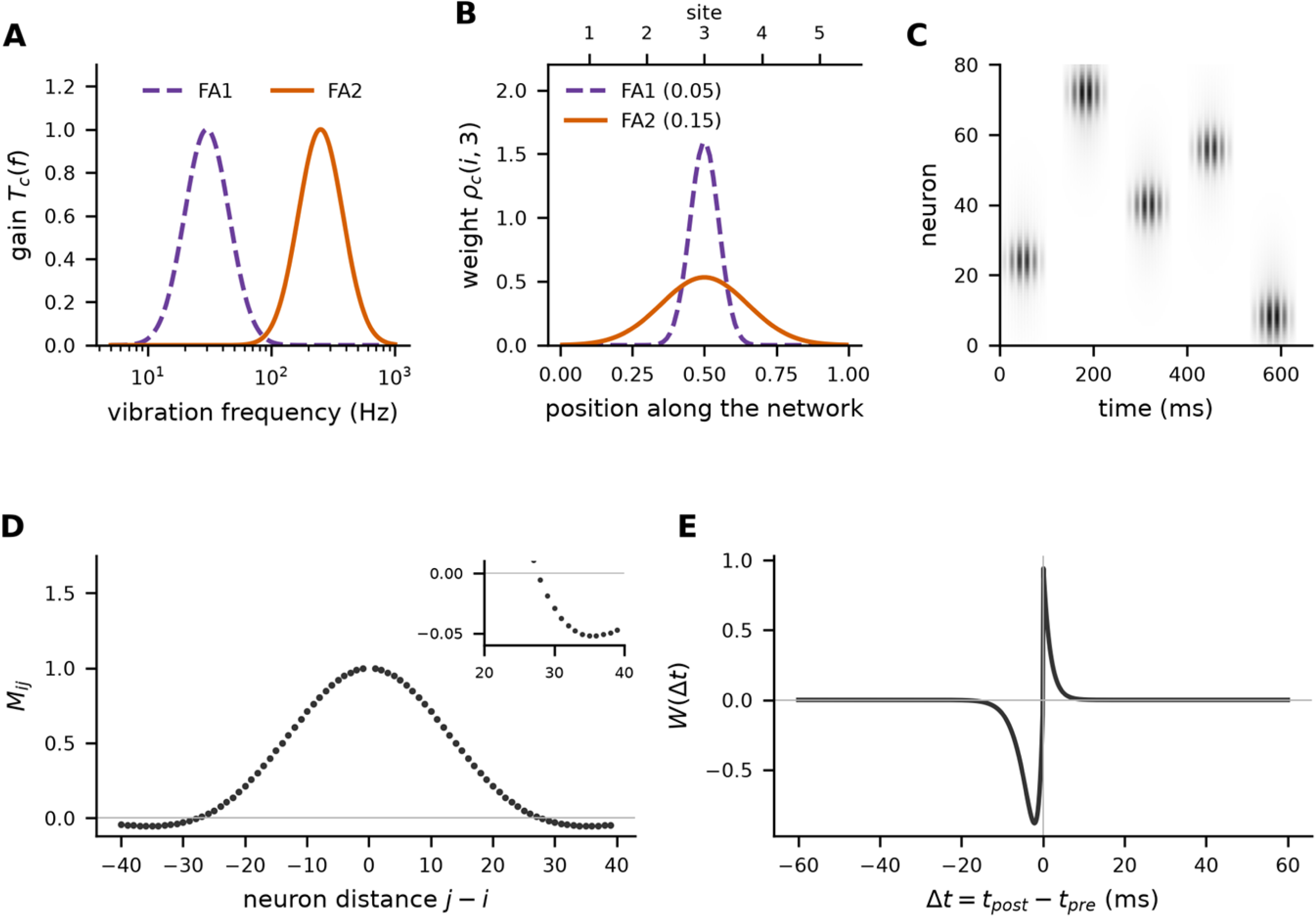
Model structure. (A) Log-Gaussian frequency tuning of the FA1 and FA2 channels, with best frequencies of 30 and 250 Hz and a bandwidth of 0.6 octaves. (B) Receptive fields of the focal (FA1) and broad (FA2) channels for site 3; ticks mark the five site centres. (C) Stimulation drive *D_i_(t)* to all 80 neurons during one CR cycle of dual-band stimulation; darker shading indicates stronger drive. (D) Mexican-hat connectivity *M*_ij_ as a function of neuron distance; the inset enlarges the inhibitory range. (E) STDP kernel *W(Δ t)*.

The dual-band burst produced the lowest coupling of the eleven patterns and improved a clinical-style protocol, at one third of the carrier’s drive power in model units. Its advantage did not follow from dose or envelope, and it rested on two properties that could be separated by control conditions: the 30 Hz component was phase-locked to the burst envelope, which left the stimulated neurons silent for 50 to 120 ms after every burst, and its focal receptive field confined part of the drive to the stimulated sector. Shifting the phase of the 30 Hz component removed the advantage without changing dose, spectrum or timing, and a second 250 Hz component delivered through a focal field reproduced it without any 30 Hz content.

## Results

### Dual-band bursts produce the lowest synaptic coupling of eleven vibrotactile CR patterns

At the end of the baseline period the network was strongly synchronized (*R* = 0.8250, SD 0.0736) and its coupling had risen to *C*_av_ = 0.1626 (SD 0.0233), the pathological state that anti-kindling reverses. Staggered delivery was necessary to reverse it. Simultaneous stimulation of all five sites increased synchrony and left coupling above baseline, whereas every staggered burst pattern drove coupling to negative values and maintained desynchronization in all networks (Table 1, Fig 3A, S1 Fig, S2 Fig and S3 Fig).

**Fig 3.**
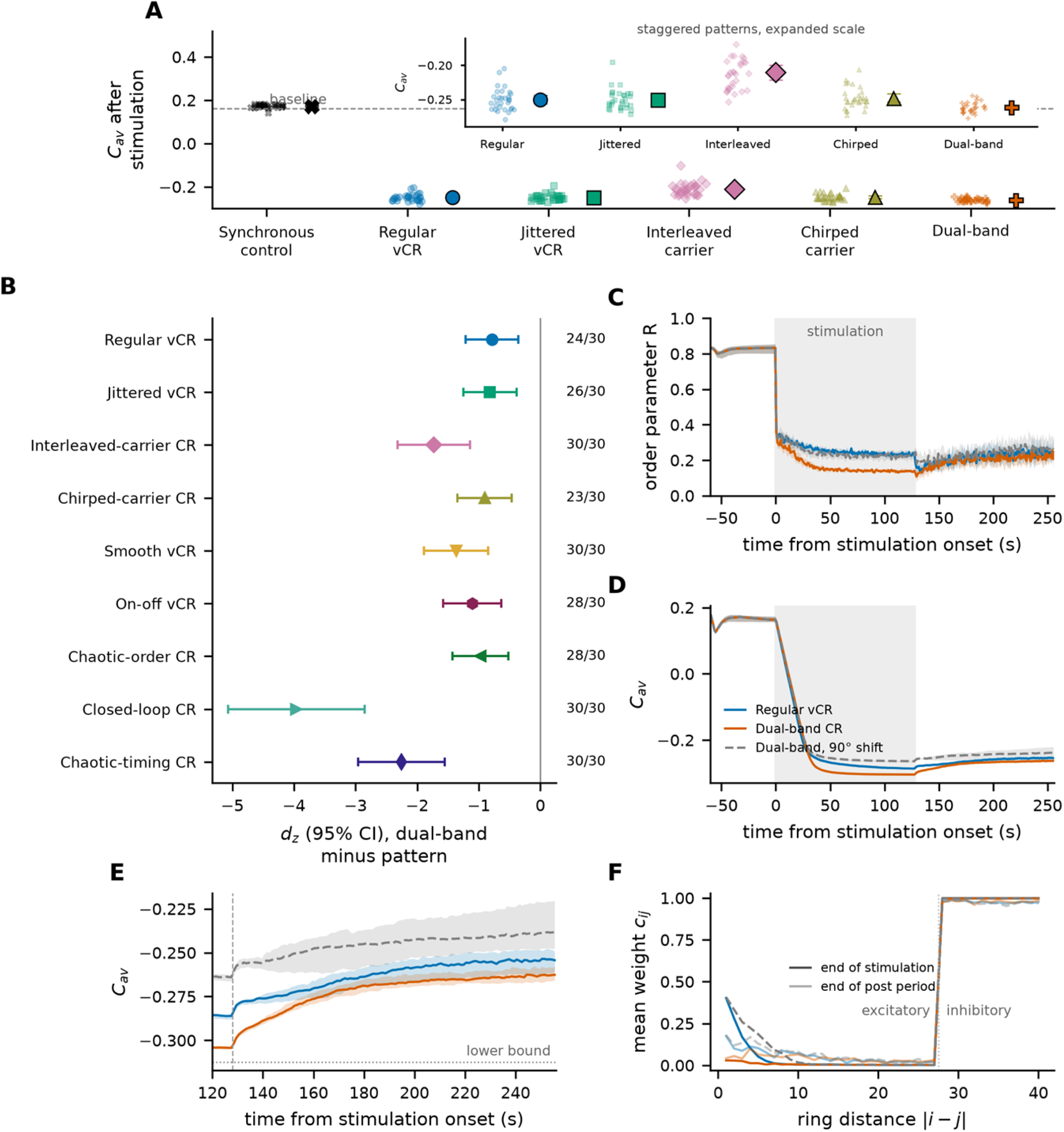
Dual-band bursts produce the lowest post-stimulation coupling, and the difference arises during stimulation. (A) *C*_av,post_ for the six patterns presented in the main text. Small symbols, individual networks (n = 30 per pattern); large symbols with error bars, mean and 95% confidence interval; dashed line, baseline mean. The inset shows the five staggered patterns on an expanded scale. (B) Paired effect sizes (*d_z_*, 95% CI) of the dual-band burst relative to each of the other patterns, with the number of networks in which coupling was lower; all ten comparisons, including the synchronous control, which lies outside the plotted range at *d_z_* = −36.1, remain significant after Holm correction at α = 0.05. (C) Order parameter and (D) mean signed coupling from the end of baseline through stimulation (shaded) and the post-stimulation period, on an independent set of 30 networks: median (line) and interquartile range (shading). (E) Coupling from the end of stimulation onwards, on an expanded scale; dotted line, lower bound of *C*_av_; dashed line, end of stimulation. (F) Mean synaptic weight as a function of ring distance in one network, at the end of stimulation (dark) and at the end of the post-stimulation period (light); dotted line, boundary between excitatory and inhibitory connections.

**Table 1.** Outcome measures for all eleven stimulation patterns.

| **Pattern** | ***R*<sub>on</sub>** | ***R*<sub>post</sub>** | ***C*<sub>av,post</sub>** | **\|M\|-weighted *C*<sub>av,post</sub>** | **Maintained** | **Drive power** |
|---|---|---|---|---|---|---|
| Synchronous multi-site control | 0.8966 (0.0097) | 0.8532 (0.0297) | 0.1731 (0.0079) | 0.4630 (0.0058) | 0/30 | 1.00 |
| Regular vCR (250 Hz carrier) | 0.2327 (0.0111) | 0.2476 (0.0590) | −0.2500 (0.0166) | 0.0570 (0.0183) | 30/30 | 1.00 |
| Jittered vCR | 0.2682 (0.0160) | 0.2530 (0.0484) | −0.2503 (0.0151) | 0.0565 (0.0174) | 30/30 | 1.00 |
| Interleaved-carrier CR | 0.2515 (0.0135) | 0.3062 (0.0731) | −0.2104 (0.0295) | 0.1214 (0.0371) | 30/30 | 1.00 |
| Chirped-carrier CR | 0.2193 (0.0108) | 0.2534 (0.0496) | −0.2475 (0.0153) | 0.0605 (0.0188) | 30/30 | 0.86 |
| Dual-band CR | 0.1431 (0.0108) | 0.2279 (0.0303) | −0.2619 (0.0078) | 0.0319 (0.0098) | 30/30 | 0.32 |
| Smooth vCR (S) | 0.0921 (0.0116) | 0.3545 (0.0959) | −0.2027 (0.0433) | 0.1213 (0.0427) | 30/30 | 6.67 |
| On-off vCR (S) | 0.2435 (0.0253) | 0.3167 (0.1023) | −0.2122 (0.0474) | 0.1113 (0.0498) | 30/30 | 0.60 |
| Closed-loop CR (S) | 0.1851 (0.1442) | 0.2411 (0.2699) | −0.0181 (0.0615) | 0.3929 (0.0480) | 27/30 | 1.00 |
| Chaotic-order CR (S) | 0.2296 (0.0107) | 0.2586 (0.0546) | −0.2480 (0.0168) | 0.0569 (0.0185) | 30/30 | 1.00 |
| Chaotic-timing CR (S) | 0.4714 (0.1088) | 0.6359 (0.2346) | 0.0409 (0.1347) | 0.3628 (0.1089) | 15/30 | 0.12 |

Mean (SD) of 30 paired networks. Maintained: number of networks with *R*<sub>post</sub> below 90% of baseline. Drive power is normalized to regular vCR. (S), reported in S1 Text. Baseline values for all patterns: *R* 0.8250 (0.0736), *C*<sub>av</sub> 0.1626 (0.0233).

The eleven patterns differed in the coupling that remained after stimulation (Friedman test, *χ*^2^(10) = 258.2, *p* < 10⁻⁴⁸), and the dual-band burst reached the lowest value of all of them (*C*_av,post_ = −0.2619, SD 0.0078). Against the 250 Hz carrier (−0.2500, SD 0.0166) it lowered coupling by 0.0119 and did so in 24 of 30 paired networks (*d_z_* = −0.78 [−1.21, −0.36], *p* = 1.8 × 10⁻⁴), and it ranked below each of the other nine patterns after Holm correction (Fig 3B). The ranking did not depend on the data used to establish it: the dual-band burst had the lowest mean coupling on one half of the networks and outperformed every other pattern on the other half, and it ranked first in 1000 of 1000 bootstrap resamples. Weighting each connection by its anatomical strength separated the two stimuli further (|M|-weighted coupling 0.0319, SD 0.0098, against 0.0570, SD 0.0183; lower in 29 of 30; *d_z_*= −1.48 [−2.02, −0.94]). The remaining patterns that varied burst content, which alternate the two carrier frequencies between cycles or sweep the carrier within each burst, fell short of the dual-band burst, as did onset jitter applied to the carrier (Table 1). All staggered patterns reached 89% to 95% of the attainable range of *C*_av_, so the differences between them are small in absolute terms.

Time courses recorded on an independent set of 30 networks, held back from every parameter choice, located the origin of this difference (Fig 3C to 3F). Both stimuli desynchronized the network within the first seconds, after which the dual-band burst held the order parameter below the carrier for the remainder of the stimulation period (median 0.14 against 0.24; lower in all networks, *d_z_*= −6.88 [−8.73, −5.02]). Coupling fell steeply under both stimuli and reached a plateau after about 40 s, with the dual-band burst below the carrier in every network from 30 s onwards and 0.0177 lower at the end of stimulation (*d_z_*= −8.07 [−10.24, −5.91]). Neither stimulus reached the lower bound of the measure, so the difference is not an artefact of saturation. Coupling then recovered under both stimuli, and the difference narrowed to 0.0103 (−0.2612, SD 0.0077, against −0.2509, SD 0.0160; lower in 23 of 30; *d_z_* = −0.58 [−0.99, −0.18], *p* = 3.4 × 10⁻³), which reproduces the main comparison on networks that played no part in selecting any parameter. The lasting advantage therefore combines a lower coupling at the end of stimulation with less recovery afterwards. Both stimuli reshaped the weight matrix in the same way, depressing excitatory connections within about 27 neurons of the diagonal and potentiating the more distant inhibitory ones; they differed in how far the short-range excitatory weights recovered once stimulation stopped (Fig 3F, S3 Fig).

### Dose and envelope do not account for the advantage

The dual-band burst differs from the carrier in dose, envelope, oscillatory content and receptive-field coverage. Control conditions that changed one property at a time (Table 2) were compared with the dual-band burst in a single family of ten tests (Fig 4), alongside the phase variants treated in the next section.

**Fig 4.**
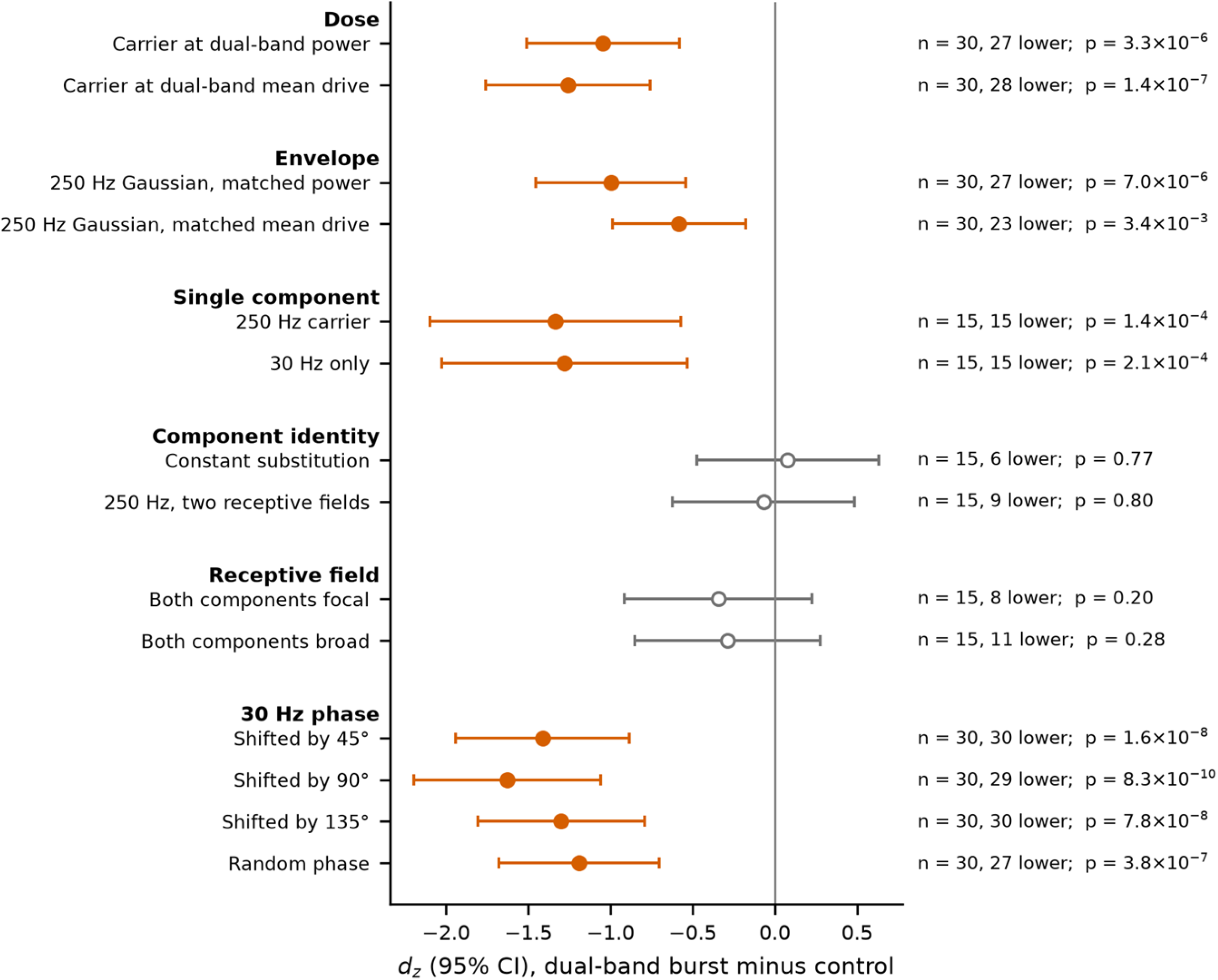
The advantage survives matched dose and envelope and depends on the phase of the 30 Hz component. Paired effect sizes (*d_z_*, 95% CI) of the dual-band burst relative to each control condition for *C*_av,post_, grouped by the property tested. The ten control conditions form one Holm family; the four phase variants, which use the schedules of the phase-locked burst, are tested separately. Filled symbols, coupling lower with the dual-band burst; open symbols, no difference. Numbers on the right give the number of networks, the number in which coupling was lower with the dual-band burst, and uncorrected *p* values.

**Table 2.** Control conditions.

| **Control condition** | **Description** | **Property tested** |
|---|---|---|
| Carrier at dual-band power | Regular vCR, amplitude 0.564 | The advantage reflects lower drive power |
| Carrier at dual-band mean drive | Regular vCR, amplitude 0.718 | The advantage reflects lower mean drive |
| 250 Hz Gaussian, matched power | Single 250 Hz component with Gaussian envelope, amplitude 0.849 | The advantage reflects the Gaussian envelope |
| 250 Hz Gaussian, matched mean drive | Single 250 Hz component with Gaussian envelope, amplitude 1.2 | The advantage reflects the Gaussian envelope |
| 250 Hz carrier only | Regular vCR, amplitude 1 | One component is sufficient |
| 30 Hz only | Regular timing, single 30 Hz rectangular component, amplitude 1 | The 30 Hz component alone is sufficient |
| Constant substitution | Dual-band burst with the 250 Hz component replaced by a constant of equal mean under the same envelope | The 250 Hz oscillation is required |
| 250 Hz, two receptive fields | Single 250 Hz rectangular component projected through the broad field and through a focal (0.05) field, amplitude 0.707 each | A second frequency is required, rather than a second spatial channel |
| Both components focal | Dual-band burst with the 250 Hz component projected through a focal (0.05) field | The focal and broad pairing is required |
| Both components broad | Dual-band burst with the 30 Hz component projected through a broad (0.15) field | The focal and broad pairing is required |
| 30 Hz shifted by 45°, 90° or 135° | Dual-band burst with the 30 Hz component shifted against the envelope | The advantage requires the 30 Hz component to be phase-locked to the envelope |
| 30 Hz at a random phase | Dual-band burst with a new phase of the 30 Hz component in every burst | The advantage requires the phase to be fixed across bursts |
| Clinical-style replica | Onset jitter and three-on, two-off cycling; 250 Hz rectangular bursts | Reference for the clinical-style protocol |
| Clinical-style dual-band | Clinical-style replica with dual-band bursts | The dual-band burst improves the clinical-style protocol |

Each condition changes one property of the dual-band burst or of the carrier.

Matching dose left the advantage intact. Reducing the amplitude of the carrier to the drive power or the mean drive of the dual-band burst, and replacing the rectangular envelope of the carrier by the Gaussian envelope of the dual-band burst at either matched dose, all produced higher coupling than the dual-band burst (dual-band lower in 23 to 28 of 30 networks, *d_z_* between −0.58 and −1.26). The Gaussian envelope on its own thus improved on the rectangular carrier at equal mean drive without reaching the dual-band burst. Neither component in isolation reproduced the effect: a single 250 Hz carrier and a single rectangular 30 Hz component both gave higher coupling in all 15 networks tested, and the 30 Hz component alone was by far the weakest of the two (*C*_av,post_ = −0.1627, SD 0.0776). All six differences survived Holm correction across the ten control comparisons (largest *p* = 3.4 × 10⁻³).

Four controls instead reached the level of the dual-band burst. Equivalence within a margin of ±0.006 was established for a burst whose 250 Hz oscillation was replaced by a constant of equal mean under the same envelope, for a burst in which a second 250 Hz component projected through a focal field took the place of the 30 Hz component, and for a burst with both components projected through the focal field (two one-sided tests, *p* = 0.005, 0.002 and 0.017). Projecting both components through the broad field gave an intermediate value for which equivalence was not established. The oscillation at 250 Hz is therefore dispensable, and a focal second component can substitute for the 30 Hz component. The two sections that follow identify the properties that these controls share with the dual-band burst.

### Phase locking of the 30 Hz component produces post-burst pauses

In the dual-band burst the 30 Hz component completes exactly three cycles under the envelope, so that its rectified waveform returns to zero at burst offset together with the envelope. Isolated neurons at the centre of a stimulated sector responded to this burst with a pause in firing, whereas the regular vCR burst was followed by uninterrupted firing (Fig 5A). During the burst the neuron fired at an elevated rate; after its last spike the final lobe of the 30 Hz component decayed to zero while the membrane recovered, leaving the potential close to its resting value, from which repetitive firing resumed only after a delay. Across the range of input currents used in the network the longest interval after the phase-locked burst averaged 83 ms (range 56 to 121 ms), against 16 ms after the regular vCR burst and an intrinsic interval of about 14 ms.

**Fig 5.**
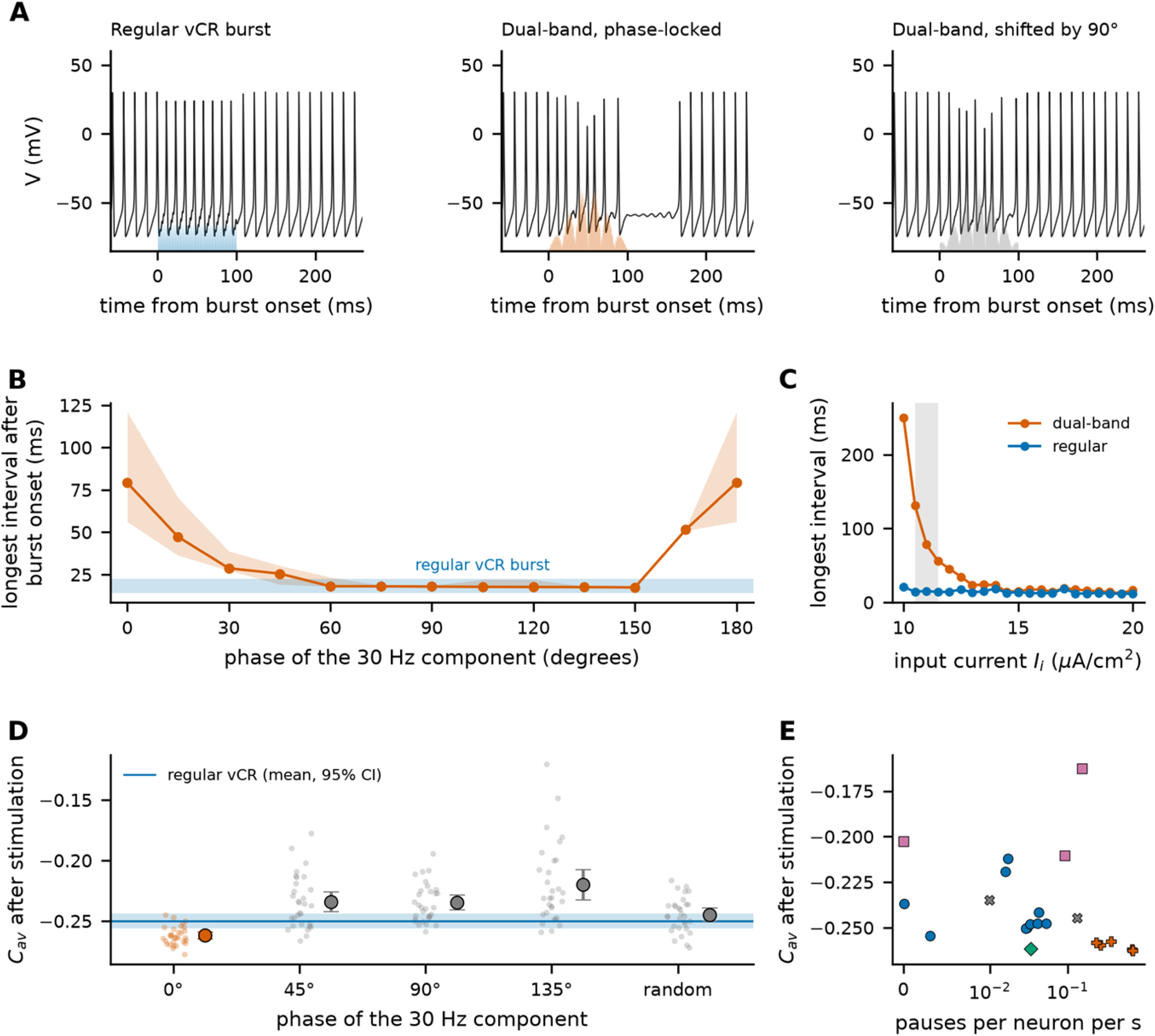
A phase-locked 30 Hz component produces post-burst pauses that carry the advantage. (A) Membrane potential of an isolated Hodgkin-Huxley neuron at a sector centre (*I_i_* = 11 µA/cm²) during and after one burst; the shaded area shows the drive *D_i_(t)* delivered to that neuron. (B) Longest inter-spike interval after burst onset as a function of the phase of the 30 Hz component: median (line) and range (shading) over the 16 input currents spanning the range used in the network; the band shows the range for the regular vCR burst. The rectified component has a period of 180°, so the values at 0° and 180° are identical. (C) Longest interval as a function of input current for both bursts; the shaded band marks the range used in the network. (D) *C*_av,post_ in the network for the phase-locked burst and for the phase-shifted variants (n = 30 networks each, same schedules); small symbols, individual networks; large symbols, mean and 95% confidence interval. (E) *C*_av,post_ against the rate of post-burst pauses during stimulation for every pattern and control. Symbols group the conditions by burst content: circles, a single broad 250 Hz component; filled crosses, dual-band bursts with a phase-locked 30 Hz component; open crosses, dual-band bursts with a shifted or randomized phase; diamond, the focal and broad 250 Hz control; squares, the remaining patterns. Three patterns that left coupling above −0.15 lie outside the plotted range.

The pause required the 30 Hz component to be aligned with the envelope (Fig 5B). It appeared when the burst ended within about 15° of a zero of the rectified component and was absent at intermediate phases, at which the burst ends on a rising or falling lobe. It also required the neurons to sit close to the onset of repetitive firing (Fig 5C): the longest interval fell from 250 ms at 10.0 µA/cm² to 78 ms at 11.0 µA/cm² and 24 ms at 13.0 µA/cm², and above 14 µA/cm² it matched the response to the regular vCR burst.

This dependence carried through to anti-kindling in the network (Fig 5D). Shifting the 30 Hz component by 90° against the envelope, while leaving site order, burst onsets, dose and spectral content untouched, raised coupling from −0.2619 (SD 0.0078) to −0.2347 (SD 0.0160), higher in 29 of 30 networks (*d_z_* = 1.63 [1.06, 2.20], *p* < 10⁻⁹), and above the level of the carrier. Shifts of 45° and 135° had the same effect, and drawing a new phase for every burst gave an intermediate value that did not differ from the carrier (*p* = 0.14). Both results held on the independent set of networks. Across conditions the rate of post-burst pauses during stimulation tracked the coupling reached afterwards (Fig 5E; Spearman *ρ* = −0.59, *p* = 2.8 × 10⁻³ over 23 conditions): every condition with more than 0.2 pauses per neuron per second ended below −0.25, and the three conditions with the most pauses were the phase-locked dual-band burst, the constant-substitution control and the clinical-style dual-band protocol.

Synchrony during stimulation followed the same pattern. The phase-locked burst gave the lowest synchrony of all conditions (*R*_on_ = 0.1431, SD 0.0108), the phase-shifted burst reached only the level of the carrier, and the random-phase burst lay between the two. The pause produced by the phase-locked burst therefore acts during stimulation, where CR theory places the decorrelation that drives anti-kindling.

### Receptive-field overlap couples the stimulated sectors

The second route to low coupling is spatial. The five stimuli were equally decorrelated at the sites themselves, but the input that each sector of the network receives is not, because the broad FA2 field spreads the input of each site over neighbouring sectors. Measured after the receptive-field projection, the input of the carrier was 1.8 times more coherent across sectors than that of the dual-band burst, and the carrier delivered 57% of each site’s drive to the sector centred on it, against 77% for the dual-band burst, whose focal component is confined to that sector (Fig 6A).

**Fig 6.**
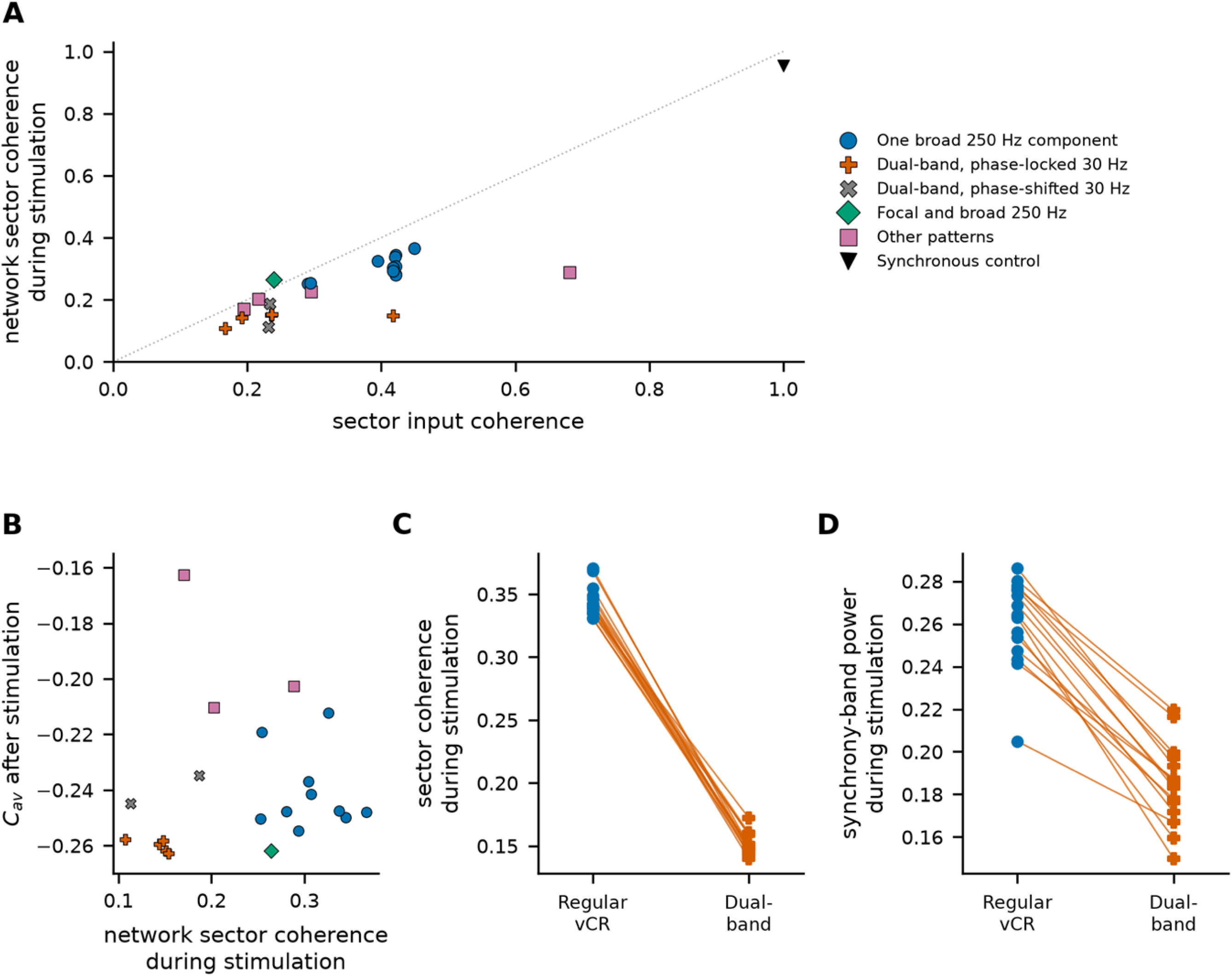
Receptive-field overlap couples the stimulated sectors, and spatial selectivity acts independently of the pause. (A) Coherence between the inputs that the five sectors receive, measured after the receptive-field projection, against the coherence of network activity between sectors during stimulation, for every pattern and control (means over networks); the dotted line marks identity. (B) *C*_av,post_ against network sector coherence during stimulation; three patterns that left coupling above −0.15 lie outside the plotted range. (C) Sector coherence and (D) synchrony-band power during stimulation for regular vCR and the dual-band burst in paired networks (n = 15).

Within the network, sector coherence during stimulation was 2.3-fold lower with the dual-band burst than with the carrier (0.1513, SD 0.0083, against 0.3440, SD 0.0123; lower in all 15 networks; *d_z_* = −11.5 [−16.1, −7.0]; Fig 6C). The network therefore adds a further factor of about 1.3 to the separation already present in the input. The dual-band burst also suppressed the intrinsic 71.5 Hz rhythm more strongly (Fig 6D and S4 Fig).

Spatial selectivity acted independently of the 30 Hz component. Replacing that component by a second 250 Hz component projected through a focal field, which carries no 30 Hz content and produces almost no pauses, gave the same coupling as the dual-band burst even though its sector coherence during stimulation remained at the level of the carrier. Conversely, the broad-field variant of the dual-band burst, whose input is as coherent across sectors as that of the carrier, still reached −0.2582 through its pauses alone. Across conditions, neither sector input coherence nor network sector coherence predicted the coupling reached afterwards (Spearman *ρ* = 0.24 and 0.29, *p* = 0.28 and 0.18; Fig 6B), which shows that decorrelation during stimulation and lasting coupling can dissociate, as reported for CR in networks with distance-dependent connectivity [11]. Each of the two properties identified here, a phase-locked low-frequency component and a focal second component, lowers the coupling that remains after stimulation, and the dual-band burst carries both.

### The dual-band burst strengthens anti-kindling within a clinical-style protocol

Substituting the dual-band burst for the 250 Hz burst of a clinical-style protocol, with onset jitter and three-on, two-off cycling unchanged, strengthened anti-kindling (Fig 7). Coupling fell by 0.0385 [0.0224, 0.0546] relative to the clinical-style replica (−0.2577, SD 0.0107, against −0.2192, SD 0.0401; lower in 26 of 30 networks; *d_z_* = −0.98 [−1.43, −0.52], *p* = 9.8 × 10⁻⁶), at one third of its drive power. The replica left a tail of networks in which little anti-kindling occurred, and the dual-band burst removed it: 28 of 30 networks reached coupling below −0.24, against 12 of 30 with the replica (Fig 7E). The |M|-weighted coupling and the synchrony measured during and after stimulation agreed after Holm correction. Duty-cycled stimulation with the dual-band burst also gave lower coupling than continuous stimulation with the carrier (*d_z_* = −0.45 [−0.84, −0.06], *p* = 0.020) at about one fifth of its drive power.

**Fig 7.**
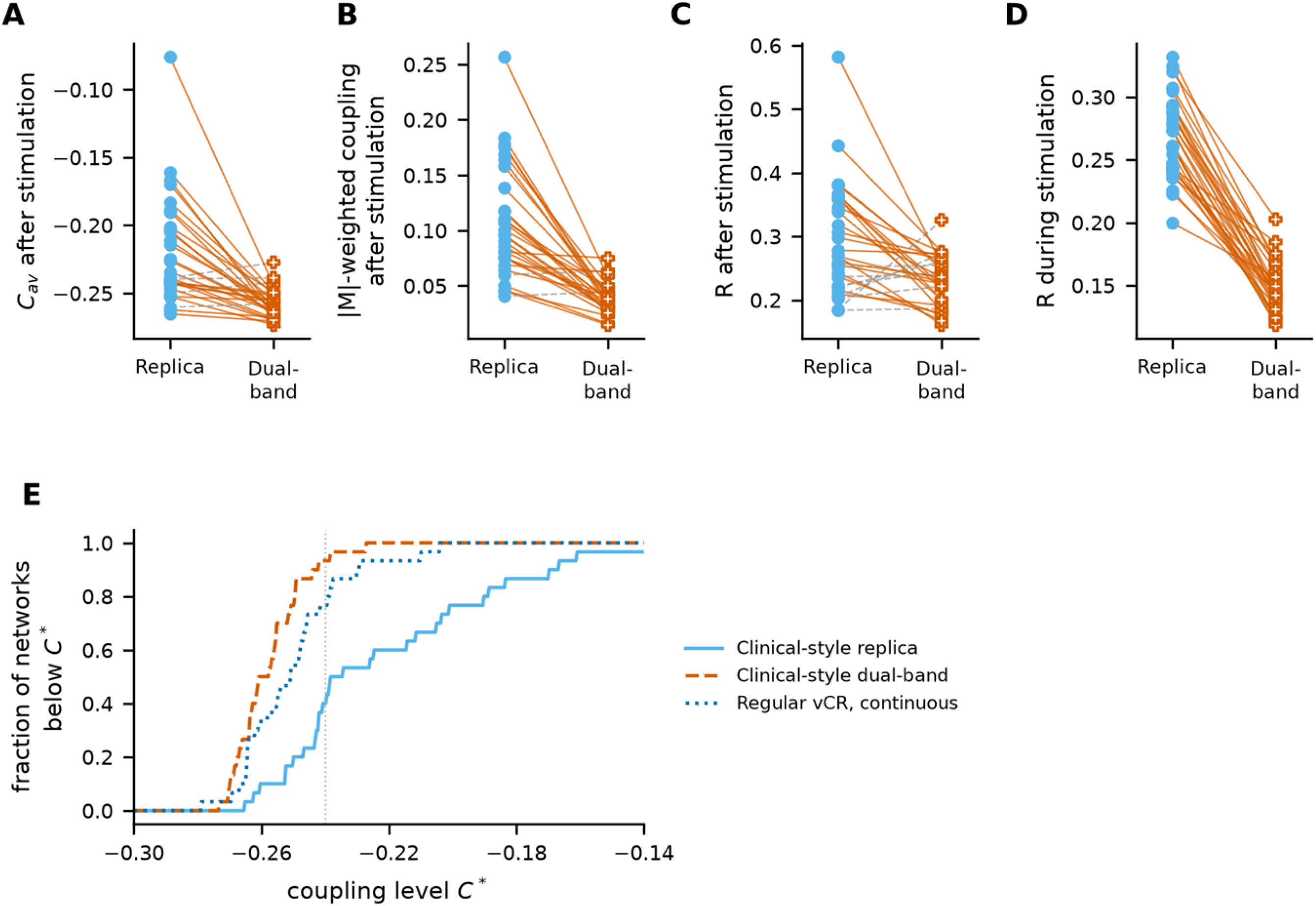
The dual-band burst strengthens anti-kindling within a clinical-style protocol. Clinical-style replica (onset jitter, three-on, two-off cycling, 250 Hz rectangular bursts) and clinical-style dual-band protocol (same timing, dual-band bursts) in 30 paired networks. (A) *C*_av,post_, (B) |M|-weighted coupling after stimulation, (C) *R*_post_ and (D) *R*_on_; solid lines mark networks with lower values under the dual-band protocol. (E) Fraction of networks reaching a coupling level *C*\* after stimulation, for the two clinical-style protocols and for continuous regular vCR; the dotted line marks *C*\* = −0.24.

The deployed device stimulates four fingertips of each hand, which gives slots of 166.7 ms and a jitter window of ±19.6 ms rather than the five sites, 133.3 ms slots and ±31 ms used above [19,22]. Repeating the comparison with four sites and the corresponding jitter gave the same result: coupling was lower with the dual-band burst by 0.0291 (−0.2536, SD 0.0132, against −0.2245, SD 0.0357; lower in 25 of 30 networks; *d_z_* = −0.79 [−1.22, −0.37], *p* = 1.5 × 10⁻⁴), as were the |M|-weighted coupling and both synchrony measures, and 27 of 30 networks reached coupling below −0.24 against 11 of 30 under the replica.

### The effect is robust to network size, model parameters and component amplitude

The direction of the effect was preserved under each of nine single-parameter changes, tested as one Holm family (Fig 8A). It remained significant after correction when the plasticity rate was halved and at both altered widths of the Mexican-hat profile, and it kept its sign with a smaller magnitude at either altered stimulation strength and at the doubled plasticity rate. The stronger desynchronization during stimulation was significant in every condition and across the calibrated range of *K*_stim_ (Fig 8C). In a network of 200 neurons the dual-band burst again gave lower coupling than the carrier (−0.2633, SD 0.0029, against −0.2596, SD 0.0055; 11 of 15 networks; *d_z_*= −0.71 [−1.33, −0.09]) and than jittered vCR, with the |M|-weighted coupling and the synchrony during stimulation agreeing.

**Fig 8.**
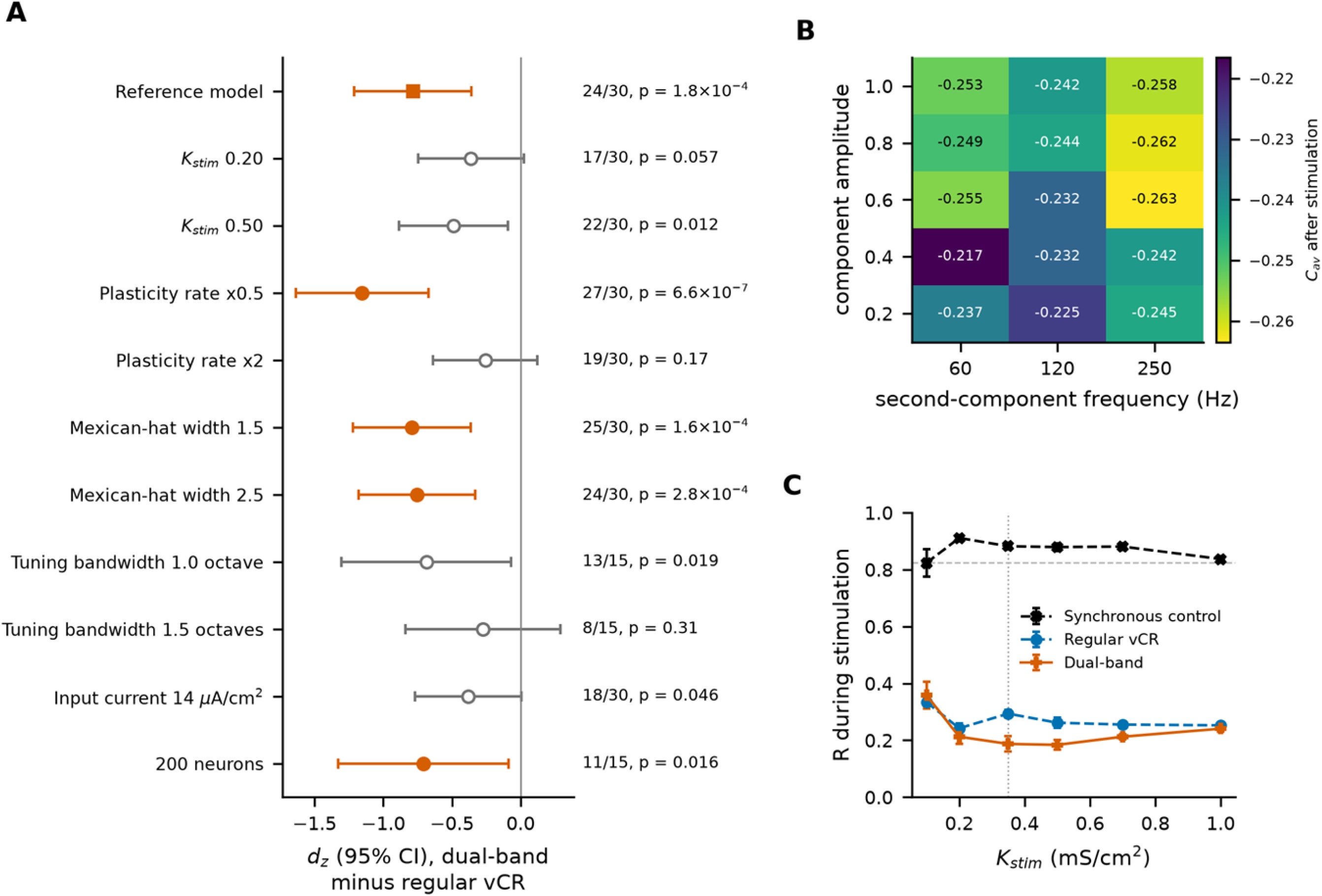
The effect is robust to network size and model parameters within a range of component amplitudes. (A) Paired effect sizes (*d_z_*, 95% CI) of the dual-band burst relative to regular vCR for *C*_av,post_ in the reference model, under single-parameter changes (n = 30, or n = 15 for the tuning bandwidth) and in a network of 200 neurons (n = 15). Filled symbols, significant after Holm correction within the family of parameter changes (α = 0.05); open symbols, same direction, not significant. (B) Mean *C*_av,post_ as a function of component amplitude and second-component frequency, with the first component fixed at 30 Hz (n = 15 per condition). (C) Order parameter during 30 s of stimulation as a function of *K*_stim_ (mean ± SD, n = 5); the dashed line marks the baseline value and the dotted line the value used.

Two of the nine changes bear directly on the two routes. Broadening the tuning of both afferent channels, so that each component also drives the other channel and the focal and broad fields are no longer separated by frequency, reduced the advantage in proportion: at a bandwidth of 1.0 octave it was smaller (*d_z_* = −0.69 [−1.30, −0.07]) and at 1.5 octaves it was no longer detectable (*d_z_*= −0.27 [−0.84, 0.29]). Raising the input current of every neuron to 14 µA/cm², which moves the network well above the onset of repetitive firing and abolishes the post-burst pause in isolated neurons, halved the advantage, from 0.0119 to 0.0052 (*d_z_* = −0.38 [−0.77, 0.01]), although the dual-band burst still desynchronized the network more strongly during stimulation. Each manipulation removes one of the two routes and weakens the effect accordingly.

The effect also depended on the amplitude of the two components (Fig 8B). With the second component at 250 Hz, amplitudes of 0.6, 0.8 and 1.0 produced lower coupling than the carrier, whereas amplitudes of 0.2 and 0.4 did not differ from it. A second component at 250 Hz gave lower coupling than one at 60 or 120 Hz at every amplitude tested (Table C in S1 Text). At a tuning bandwidth of 0.6 octaves a 60 Hz component is routed to FA1 and a 120 Hz component to FA2 at low gain, so this axis varies the strength of the broad component rather than its frequency alone.

## Discussion

A dual-band burst whose 30 Hz component is phase-locked to the burst envelope strengthened anti-kindling by vibrotactile CR in a spiking network with STDP. It produced the lowest synaptic coupling of eleven vibrotactile CR patterns, replicated on an independent set of networks, and improved a clinical-style protocol when it replaced the 250 Hz burst, while delivering about one third of the carrier’s drive power in model units. Control conditions excluded dose and envelope as explanations and identified two properties through which a burst reaches this level of coupling: a pause in firing produced by the phase-locked 30 Hz component, and a focal component that confines part of the drive to the stimulated subpopulation. The dual-band burst engages both, and either one alone is sufficient to reach the same coupling.

The first property connects burst waveform to the principle on which CR rests. CR desynchronizes by resetting the phase of each stimulated subpopulation at a different time [5]. A model of vibrotactile CR with leaky integrate-and-fire neurons assumed instead that vibratory stimuli do not reset neuronal phase and act through a rate increase during the burst [19]. Our results indicate that both descriptions can hold, depending on the waveform of the burst and on the excitability of the target neurons. When the drive returned to zero at burst offset, neurons biased just above the onset of repetitive firing settled near their resting state and resumed firing only after 50 to 120 ms, with a delay that varied with each neuron’s input current. A pause of this kind, locked to the end of every burst, is a phase reset of the stimulated sector, and it is the feature that distinguishes the phase-locked burst from the same burst with its 30 Hz component shifted by 90°. The pause depends on type 2 excitability, in which repetitive firing begins at a Hopf bifurcation and the resting state remains weakly stable or weakly unstable nearby [31]. Fast-spiking interneurons of somatosensory cortex show this behaviour, including irregular switching between firing and subthreshold oscillation close to threshold [30], whereas regular-spiking pyramidal neurons do not. Whether vibration bursts produce such pauses in the populations targeted by vibrotactile CR is therefore an experimental question, and the prediction is specific: the effect should depend on the alignment of the low-frequency component with the burst envelope, not on its presence.

The second property is spatial and follows from afferent physiology. The 250 Hz carrier recruits mainly FA2 afferents, whose receptive fields extend well beyond the stimulated fingertip [26,28]. In the model, this spread was measurable at the input: with the carrier, the drive reaching each sector was 1.8 times more coherent across sectors than with the dual-band burst, and only 57% of each site’s drive reached the sector centred on it, against 77% with the dual-band burst. Wider spatial spread of stimulation has long been known to weaken CR [29], and the sensitivity of CR to the ratio between the spatial reach of each stimulus and the spacing of the sites has been shown in networks with distance-dependent connectivity [11]. The present model places that constraint in the afferent periphery: a carrier frequency chosen for FA2 tuning also fixes the spatial resolution of the stimulation. Recruiting focal afferents restores the separation, and a second 250 Hz component delivered through a focal field achieved the same coupling without any 30 Hz content, which isolates selectivity from the pause mechanism.

Decorrelation during stimulation did not by itself predict the coupling that remained afterwards. Across 23 conditions, neither the coherence of the input that each sector received nor the coherence of network activity during stimulation correlated with *C*_av,post_, whereas the rate of post-burst pauses did. Two conditions illustrate the dissociation: the two-field control left sector coherence at the level of the carrier yet reached the coupling of the dual-band burst, and smooth vCR produced strong desynchronization during stimulation but left coupling above baseline. Corresponding dissociations have been reported for CR in plastic networks with spatially dependent connections, where acute desynchronization does not imply a long-term effect [11], and for the dependence of the lasting effect on stimulus shape and intra-burst structure [33]. Measures taken during stimulation are therefore useful as mechanistic correlates rather than as surrogates for the outcome.

For clinical application, the results point to a change in the drive signal of an existing device rather than in its hardware or its schedule. Vibrotactile CR is attractive precisely because it is non-invasive, and any gain in efficacy that costs nothing in invasiveness is worth pursuing, given that the lasting benefit demonstrated for electrical CR in parkinsonian primates and patients [12–14] currently requires an implanted electrode. Within a clinical-style protocol, substituting the dual-band burst for the 250 Hz burst lowered coupling by three times the difference obtained with continuous stimulation and removed a tail of networks in which the replica produced little anti-kindling, which is the behaviour that matters for a protocol delivered at a fixed dose. The amplitudes in the model are expressed in model units, and the drive power is not a measure of vibration energy at the skin. Detection thresholds at the fingertip are about 0.13 to 0.17 m/s² at 31.5 Hz and about 0.47 to 1.72 m/s² at 250 Hz, which corresponds to roughly 3.3 to 4.3 µm against 0.2 to 0.7 µm of displacement [34], so a 30 Hz component matched in sensation level requires several times the displacement of the 250 Hz carrier with the same contactor. Any statement about device power therefore requires calibration in physical units, and the present results support a claim about burst composition rather than about energy. Two further points bear on a trial: the lower-frequency component dominates the percept of a superimposed dual-frequency vibration [35], which matters for tolerability and for blinding against a 250 Hz comparator, and 30 Hz lies at the upper edge of the beta band, so interactions of the added component with sensorimotor beta activity, of which a reduction has been reported as a correlate of vibrotactile CR [19], should be examined. The intrinsic rhythm of the present network lies at 71.5 Hz and does not represent beta activity.

Several limitations bound these conclusions. The post-burst pause is a property of neurons with type 2 excitability biased close to the onset of repetitive firing, and the magnitude of the effect depends on that bias: raising the input current to 14 µA/cm² abolished the pause in isolated neurons and halved the advantage in the network, leaving the part that spatial selectivity contributes. In a model with type 1 excitability, in which firing begins at an arbitrarily low rate, this route would not be available, and the effect would rest on selectivity alone. The afferent model represents frequency tuning as a static gain for each component and includes two of the four mechanoreceptor channels of the human hand [28]; it does not include afferent spiking, adaptation or the temporal coding of vibration frequency by afferent spike timing [32]. The tuning curves of the two channels barely overlap at a bandwidth of 0.6 octaves, so that each component drives one channel, and broadening them reduces the effect; at the amplitudes used clinically, a 250 Hz carrier probably also recruits some FA1 afferents. The projection from the fingertips to the network is a single Gaussian mapping onto one somatotopic ring, without the relay stages of the somatosensory pathway, and the receptive fields do not wrap around the ring, so that the two outer sites deliver a larger share of their drive to their own sector than the three inner ones. The clinical-style protocol reproduces the timing of the deployed device but not its bilateral delivery, and each simulation covers a single 128 s session rather than months of daily use, with STDP as the only form of plasticity. This matters because the therapeutic effect of CR accumulates across sessions and depends on the daily dose, both in parkinsonian primates [13] and in patients [14,19], so a single session measures the per-session increment rather than the attainable benefit. Models with spiking afferents, multi-stage pathways and repeated sessions will be needed to establish how these results transfer to the clinical setting.

The results lead to three predictions that can be tested. First, vibrotactile CR with a dual-band burst should produce greater clinical benefit than the 250 Hz carrier at equal sensation level, and the benefit should require the low-frequency component to be phase-locked to the burst envelope, so that a burst with a randomized component phase should perform like the carrier. Second, a 250 Hz burst delivered through a focal and a broad contact area at each digit should reproduce the benefit without any 30 Hz stimulation. Third, during dual-band stimulation the cortical representations of the stimulated digits should be less coherent with one another than during carrier stimulation, as measurable with electroencephalography or magnetoencephalography, even though the vibration inputs to the digits are equally decorrelated at the fingertips; the size of this difference is predicted by the sector-level input coherence computed here. Together these predictions provide a route from the present simulations to bench testing of dual-band actuators and to clinical evaluation.

## Materials and methods

### Neuron and network model

The network follows the ring network of Manos et al. [10] and consists of *N* = 80 conductance-based Hodgkin-Huxley neurons [36]; the main comparison was repeated with *N* = 200. Voltages are given in mV, time in ms, currents in µA/cm² and conductances in mS/cm². The membrane potential of neuron *i* obeys

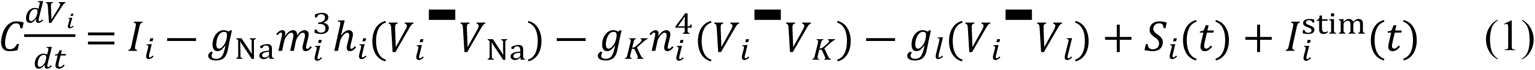

with the standard gating kinetics for *m_i_*, *h_i_* and *n_i_* [36], *C* = 1 µF/cm², *g*_Na_ = 120, *g_K_* = 36 and *g_l_* = 0.3 mS/cm², and *V*_Na_ = 50, *V_K_* = −77 and *V_l_* = −54.4 mV. The constant input current *I_i_* was drawn independently for each neuron from a uniform distribution on [10.55, 11.45] µA/cm², so that each neuron fires tonically at approximately 70 Hz with a slightly different intrinsic rate. This range lies just above the Hopf bifurcation at which the resting state of the Hodgkin-Huxley model loses stability, at about 9.8 µA/cm² [31], so that the model neurons have type 2 excitability [30]. Neurons are equally spaced on a ring of length *d*_0_ = 10 (dimensionless units). Anatomical

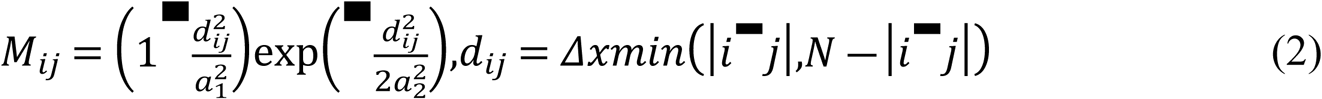

with *a*_1_ = 3.5, *a*_2_ = 2.0, *M*_ii_ = 0 and neighbour spacing *Δ x = d*_0_*/(N−1)* [10]. *M*_ij_ is excitatory for up to approximately 27 neighbours on each side and weakly inhibitory beyond, with an inhibitory minimum of approximately −0.05 relative to an excitatory maximum of 1 (Fig 2D). The synaptic input to neuron *i* is

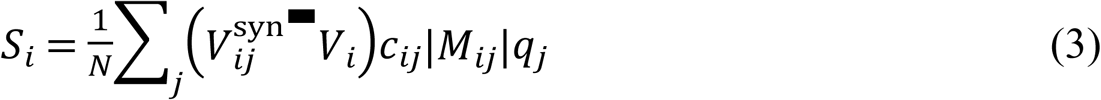

where the synaptic reversal potential *V*_ij_^*syn*^ is +20 mV for *M* > 0 and −40 mV for *M* < 0, so that the sign of *M*_ij_ determines whether a connection is excitatory or inhibitory. The synaptic gating variable *q_j_*is driven continuously by the presynaptic membrane potential,

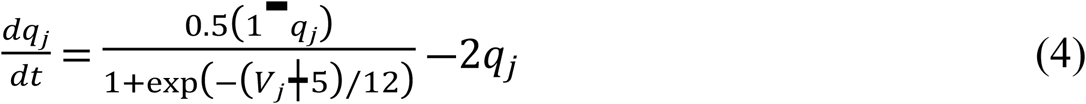

The plastic weights *c*_ij_ (from neuron *j* to neuron *i*) are dimensionless, bounded to [0, 1], and scale a maximal synaptic conductance of 1 mS/cm².

### Spike-timing-dependent plasticity

Spike times were defined as upward crossings of 0 mV by the membrane potential; the same spike times were used for the plasticity rule and for the estimation of neuronal phases. Whenever neuron *i* fires at time *t*, all weights onto and from neuron *i* are updated using the most recent spike time *t_j_*of each partner neuron:

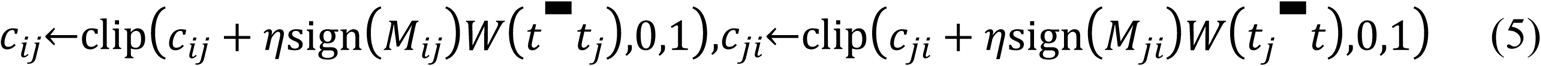

so that in both cases the argument of *W* is *Δ t = t*_post_ *− t*_pre_. The plasticity kernel is

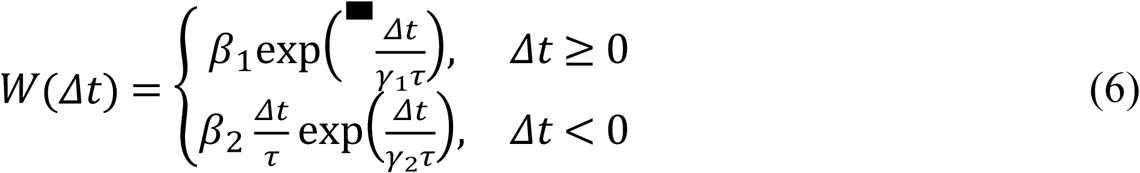

with *β*_1_ = 1, *β*_2_ = 16, *γ*_1_ = 0.12, *γ*_2_ = 0.15, *τ* = 14 ms and learning rate *η* = 0.002. Because the update is multiplied by *sign(M*_ij_), excitatory connections are potentiated by causal and depressed by anti-causal spike pairs, and inhibitory connections are modified in the opposite direction. The depressing lobe of the kernel has three times the area of the potentiating lobe (*β_2_γ*_2_^2^*τ* = 5.04 ms against *β*_1_*γ*_1_*τ* = 1.68 ms; Fig 2E), so that uncorrelated firing weakens excitatory and strengthens inhibitory connections, which is the basis of anti-kindling.

### Afferent model and stimulation current

Each vibration burst is converted into a stimulation current by two afferent channels, *c* ∈ {FA1, FA2}, each characterized by a frequency tuning curve and a receptive field on the network (Fig 2A and 2B). Both channels project onto the same neuronal population, reflecting the finding that most neurons in areas 3b and 1 of primary somatosensory cortex receive input from both FA1 and FA2 afferents [37] and that inputs from the two channels also converge in secondary somatosensory cortex [38]. Frequency tuning is Gaussian in log-frequency,

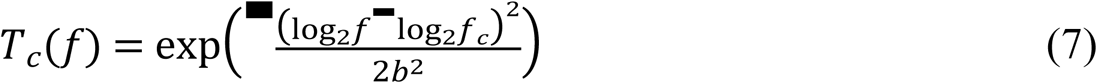

with best frequencies *f*_FA1_ = 30 Hz and *f*_FA2_ = 250 Hz and a bandwidth *b* = 0.6 octaves for both channels. With this bandwidth, a 30 Hz component drives FA1 and a 250 Hz component drives FA2 (*T*_FA1_(250 Hz) ≈ 10^−6^); broader tuning, in which each component also recruits the other channel, was examined with *b* = 1.0 and 1.5 octaves.

Five stimulation sites are located at positions *y_k_ = (k + 1/2)/5*, *k* = 0 to 4, and neuron *i* is located at *x_i_ = (i + 1/2)/N* on a unit coordinate along the ring. Each channel projects each site onto the network through a normalized Gaussian receptive field,

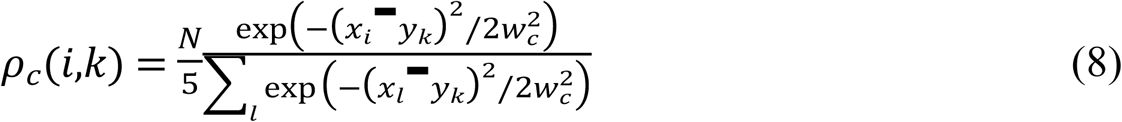

with widths *w*_FA1_ = 0.05 (focal) and *w*_FA2_ = 0.15 (broad), a threefold ratio that reflects the difference in receptive-field extent between the two afferent classes [26]. The normalization ensures that each site delivers the same total input to the network irrespective of receptive-field width. Site-to-neuron distances are measured along the map without wrapping around the ring, because the five sites represent digits arranged in a row on the hand, in which the first and last digits are not neighbours; the ring topology of the connectivity serves only to avoid boundary effects in the network. As a consequence, the broad field of the two outer sites delivers 66% of its input to the sector centred on the site, against 50% to 51% for the three inner sites; the focal field delivers 96% to 98%.

Each burst *e* is defined by its site *k_e_*, onset *t_e_*, duration *T_b_* = 100 ms and envelope *ε_e_(t)*, and contains one or two components *m* with frequency *f*_e,m_*(t)*, amplitude *κ*_e,m_ and initial phase *φ*_e,m_. The envelope is either rectangular or Gaussian, centred in the burst with a standard deviation of 25 ms. Each component is full-wave rectified, since afferents respond to skin displacement in both directions, and components are rectified separately before they are summed. The input from channel *c* at site *k* and the resulting drive to neuron *i* are

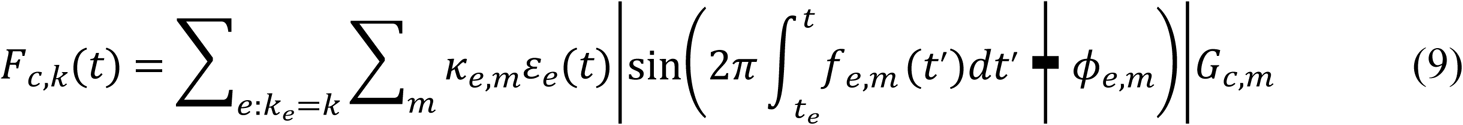

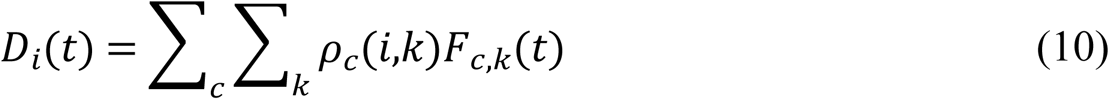

where the routing gain *G*_c,m_ equals the tuning *T_c_(f*_e,m_) unless a control condition assigns a component to a chosen receptive field. The drive enters the membrane equation as a conductance-based excitatory current,

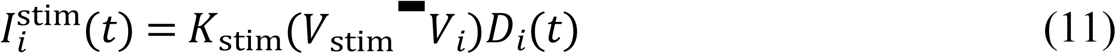

with *V*_stim_ = 20 mV and stimulation strength *K*_stim_ = 0.35 mS/cm². *K*_stim_ was calibrated with 30 s of stimulation in five networks against two criteria: the synchronous multi-site control should keep the order parameter during stimulation at or above its baseline value (0.82), and staggered stimulation with the carrier and with the dual-band burst should lower it below 0.30. Both criteria were met for every value from 0.20 to 1.00 mS/cm², and not at 0.10 mS/cm² (Table B in S1 Text); 0.35 mS/cm² lies within this range.

### Stimulation timing and the dual-band burst

All burst-based patterns share a common CR timing scheme unless stated otherwise (Fig 1C). Stimulation is organized in CR cycles of 667 ms (CR rate 1.5 Hz). At the start of each cycle the five sites are arranged in a new random order and stimulated in successive 133.3 ms slots, each site receiving one 100 ms burst per cycle.

The dual-band burst retains this timing and modifies only the content of each burst (Fig 1A and 1B). Each burst contains a 30 Hz and a 250 Hz component with identical onset and an identical Gaussian envelope, each with amplitude *κ* = 0.6. Both components start at phase zero at burst onset, so that the 30 Hz component completes exactly three cycles and its rectified waveform returns to zero at burst offset together with the envelope; we refer to this alignment as phase locking. In the phase-shifted variants, the 30 Hz component starts at a phase of 45°, 90° or 135°, or at a phase drawn uniformly for every burst; because rectification gives the component a period of 180°, these shifts span its full range. The phase-shifted variants used the same stimulation schedules as the phase-locked burst, so that they differed from it only in the phase of the 30 Hz component.

The clinical-style protocol adds two rules to the same scheme (Fig 1D): each burst onset is shifted by a random jitter, and stimulation is delivered for three CR cycles followed by two cycles without stimulation [19]. With five sites, the jitter was drawn uniformly from ±31 ms (±23.5% of the slot). With four sites, as in the deployed device, sites were stimulated in 166.7 ms slots and the jitter was drawn from ±*J T*_CR_*/(2N_s_*) = ±19.6 ms, with *J* = 23.5%, CR period *T*_CR_ and *N_s_* = 4 sites [19,22]. The clinical-style replica applies these rules to the 250 Hz rectangular burst, and the clinical-style dual-band protocol applies them to the dual-band burst.

### Stimulation patterns and control conditions

Eleven stimulation patterns were compared (Table 3). They comprise a synchronous multi-site control, the clinically used 250 Hz carrier (regular vCR), timing variants of regular vCR and patterns that differ in the content of the bursts. Six patterns are presented in the main text; the remaining five, which modify the delivery mode (continuous stimulation, duty cycling, closed-loop gating) or use chaotic scheduling, are reported in S1 Text, and all eleven were included in every statistical comparison. Single-carrier patterns use a 250 Hz carrier with a rectangular envelope and amplitude 1 unless stated otherwise. In chirped-carrier CR, the carrier frequency rises linearly from 180 to 320 Hz within each burst; in interleaved-carrier CR, the carrier alternates between 30 and 250 Hz from one CR cycle to the next.

**Table 3.** Stimulation patterns.

| Pattern | Stimulus parameters | Property tested | Source | Drive power |
|---|---|---|---|---|
| Synchronous multi-site control | All five sites stimulated simultaneously in each cycle; 250 Hz, rectangular | Desynchronization requires staggered delivery | [19] | 1.00 |
| Regular vCR | Random site order in each cycle; 250 Hz, rectangular | Reference: clinical carrier with CR timing | [10,15] | 1.00 |
| Jittered vCR | Regular vCR with random burst onset jitter (±23.5% of the slot) | Timing jitter strengthens anti-kindling | [22] | 1.00 |
| Interleaved-carrier CR | Regular timing; carrier alternates between 30 and 250 Hz from cycle to cycle | Recruiting both channels in alternate cycles strengthens anti-kindling | This study | 1.00 |
| Chirped-carrier CR | Regular timing; carrier swept from 180 to 320 Hz within each burst | A frequency-modulated carrier strengthens anti-kindling | This study | 0.86 |
| Dual-band CR | Regular timing; 30 and 250 Hz superimposed, Gaussian envelope, amplitude 0.6 each | Recruiting both channels within every burst strengthens anti-kindling | This study | 0.32 |
| Smooth vCR (S) | Continuous rectified 30 Hz sinusoid at each site, phase-shifted by *2π k/5* between sites | Continuous phase-shifted stimulation is as effective as bursts | [15,39] | 6.67 |
| On-off vCR (S) | Regular vCR, three cycles on, two off | Duty cycling preserves anti-kindling | [19] | 0.60 |
| Closed-loop CR (S) | Regular vCR suspended while the 500 ms moving average of *R* is below 0.5 | Demand-controlled stimulation preserves anti-kindling | [5] | 1.00 (scheduled) |
| Chaotic-order CR (S) | Site order determined by a logistic map (*r* = 3.9) | Chaotic site order strengthens anti-kindling | This study | 1.00 |
| Chaotic-timing CR (S) | Cycle duration (0.4 to 0.9 s) and carrier (100 to 300 Hz) determined by logistic maps; bursts of ten carrier cycles, Gaussian envelope | Chaotic timing strengthens anti-kindling | This study | 0.12 |

Drive power is given in model units relative to regular vCR. (S), reported in S1 Text.

Control conditions were designed to identify which property of the dual-band burst accounts for its effect (Table 2). Each control changes a single property of the dual-band burst or of the carrier and was compared with the dual-band burst in paired simulations.

### Stimulation dose

Two dose measures were computed from the stimulus waveforms: the model drive power, defined as the time- and site-averaged sum of squared channel inputs *Σ_c_ F*_c,k_*(t)*^2^, and the mean drive, defined as the time- and neuron-averaged *D_i_(t)*. Both are expressed in model units relative to regular vCR, and the drive power is not a calibrated measure of vibration energy at the skin. Relative to regular vCR, the dual-band burst delivers 0.32 of the drive power and 0.72 of the mean drive. The doses of all patterns and controls are listed in Table A in S1 Text.

### Simulation protocol and outcome measures

Each simulation comprised 2 s of equilibration with plasticity disabled, a 60 s baseline period with plasticity enabled and no stimulation, during which the network synchronized and its coupling strengthened, 128 s of stimulation and a 128 s post-stimulation period with plasticity enabled. Equations were integrated with the forward Euler method at a time step of 0.01 ms; the stimulus waveform was updated every 0.1 ms. Initial membrane potentials were drawn uniformly from [−65, 5] mV, gating variables uniformly from [0, 1] and initial weights from a normal distribution with mean 0.5 and SD 0.01. Each simulation used an independently drawn network (input currents, initial weights and initial state). Within a comparison, all patterns were applied to the same set of networks, so that comparisons are paired.

#### Synaptic coupling

The primary outcome was the mean signed synaptic weight

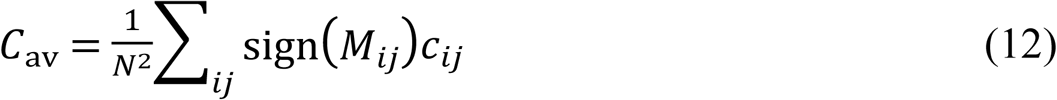

averaged over the last 5 s of the post-stimulation period (*C*_av,post_). Lower values indicate stronger anti-kindling; in this network *C*_av_ is bounded to the interval [−0.3125, 0.675]. The same average over the last 5 s of stimulation (*C*_av,stim_) measures the coupling reached during stimulation, and the difference between the two measures how much coupling recovers afterwards. Because inhibitory connections are numerous but anatomically weak, an |M|-weighted coupling, *Σ*_ij_*sign(M*_ij_*)|M*_ij_*|c*_ij_ */ Σ*_ij_*|M*_ij_|, was also computed.

#### Synchrony

Network synchrony was quantified with the Kuramoto order parameter. The phase of each neuron was estimated from its last completed inter-spike interval, *φ_i_(t) = 2π(t − t_i_*^(1)^*)/(t_i_*^(1)^ *− t_i_*^(2)^), where *t_i_*^(1)^ and *t_i_*^(2)^ are the last and second-to-last spike times, and

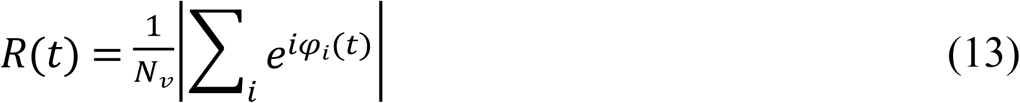

where *N_v_* is the number of neurons with at least two spikes. This estimate uses only past spikes and can therefore be computed during the simulation, as required for closed-loop gating. *R*_on_ and *R*_post_ denote the mean of *R* over the last 5 s of stimulation and of the post-stimulation period. Desynchronization was considered maintained when *R*_post_ was below 90% of the baseline value. *Network coherence and post-burst pauses.* The ring was divided into five sectors of 16 neurons, each centred on one stimulation site, and spike counts were computed in 1 ms bins. Sector coherence is the magnitude-squared coherence between sector spike counts (Welch’s method as implemented in SciPy [40], 2.048 s Hann windows, 50% overlap), averaged over the ten sector pairs and over the 0.5 to 20 Hz band, which contains the CR rate and its harmonics. Synchrony-band power is the mean power spectral density of the mean-subtracted population spike count (spikes per 1 ms bin) between 55 and 85 Hz, the band of the intrinsic network rhythm, in counts² per Hz. Post-burst pauses were counted as inter-spike intervals longer than 40 ms, more than twice the longest intrinsic interval. These measures were computed in 20 s windows at the end of baseline, at the start of stimulation and at the start of the post-stimulation period.

#### Input measures

Site input coherence was computed in the same way from the input at each site summed over channels, *Σ_c_ F*_c,k_*(t)*, and sector input coherence from the drive summed over the neurons of each sector, *Σ*_i ∈ s_ *D_i_(t)*, so that it describes the input as the network receives it after the receptive-field projection. The own-sector share is the fraction of the drive delivered from each site that reaches the sector centred on that site, averaged over sites. Input measures were averaged over three independent 20 s stimulation schedules.

#### Single neurons

To isolate the response of a neuron to one burst, isolated Hodgkin-Huxley neurons (*S_i_* = 0) at the centre of the middle sector received a single burst at that sector’s site through both receptive fields, with input currents spanning the range used in the network. The longest inter-spike interval after burst onset was recorded for the regular vCR burst and for the dual-band burst with the 30 Hz phase varied from 0° to 165° in steps of 15°.

### Statistical analysis

All comparisons were paired and two-sided, and the threshold for significance was α = 0.05, applied to the corrected *p* values within each family of tests. The number of networks per condition, 30 or 15 where stated, was fixed before the simulations were run; with 30 paired networks a two-sided paired t-test at α = 0.05 has 80% power to detect *d_z_* = 0.53, and with 15 networks *d_z_* = 0.78. Differences were tested with paired t-tests and, as a check that does not assume normally distributed paired differences, with Wilcoxon signed-rank tests, both as implemented in SciPy [40]; test statistics, degrees of freedom and both *p* values for every comparison reported in the text are listed in Table E in S1 Text. Effect sizes are reported as Cohen’s *d_z_* (mean paired difference divided by the standard deviation of the differences) with 95% confidence intervals, and differences in natural units as the mean paired difference. Negative values indicate lower coupling or synchrony with the dual-band burst. The number of simulations in which the dual-band burst gave the lower value is reported alongside. The eleven patterns were compared with a Friedman test, blocked by network, followed by paired comparisons of the dual-band burst against each of the other ten patterns. Holm’s procedure [41] was applied within each family of tests: the ten pattern comparisons, all control comparisons together, and the parameter changes. Equivalence of a control with the dual-band burst was assessed with two one-sided paired t-tests [42] at a margin of ±0.006, half of the difference in *C*_av,post_ between the dual-band burst and the carrier in the main comparison.

To check that the ranking did not depend on selection from the same data, the pattern with the lowest mean *C*_av,post_ was selected on one half of the networks and tested on the other, and the ranking by mean *C*_av,post_ was recomputed in 1000 bootstrap resamples of networks. The comparison with the carrier was repeated on an independent set of 30 networks that played no part in any parameter choice. Each study drew its own stimulation schedules, so that comparisons repeated on the same networks in different studies differ only in the random site order and indicate the variability due to the schedule. Means and standard deviations are reported to four decimal places.

### Software

Simulations were implemented in Python 3.12 with NumPy 2.4.6, Numba 0.64.0, SciPy 1.17.1, pandas 2.3.3, joblib 1.4.2 and matplotlib 3.11.0. Random seeds and the allocation of networks to analyses are listed in S1 Text.

## Data Availability

All materials are available at https://github.com/shrpatki/vcr-dualband-anti-kindling and we have archived our code on Zenodo (DOI: 10.5281/zenodo.23746222).

## Use of AI-assisted tools

The authors used Claude (Anthropic) to assist with preparing simulation code, preparing code to generate figures, and drafting and revising the manuscript text. All equations, parameter values and reported results were verified by the authors against the simulation code and output files.

## Acknowledgments

The authors would like to express sincere gratitude to Pranmya Rathaparkhi for his valuable guidance and support. The authors also acknowledge the Indian Institute of Technology Madras (IIT Madras) for providing the institutional support and resources that contributed to this work.

## Supporting information

**S1 Text. Supplementary methods, patterns, tables and test statistics.** Supplementary stimulation patterns, random seeds and the allocation of networks to analyses, numerical details, and Tables A to E (stimulation dose, calibration of the stimulation strength, dependence on component amplitude and second-component frequency, parameter changes with 15 networks per condition, and test statistics for the comparisons reported in the main text).

**S1 Fig.**
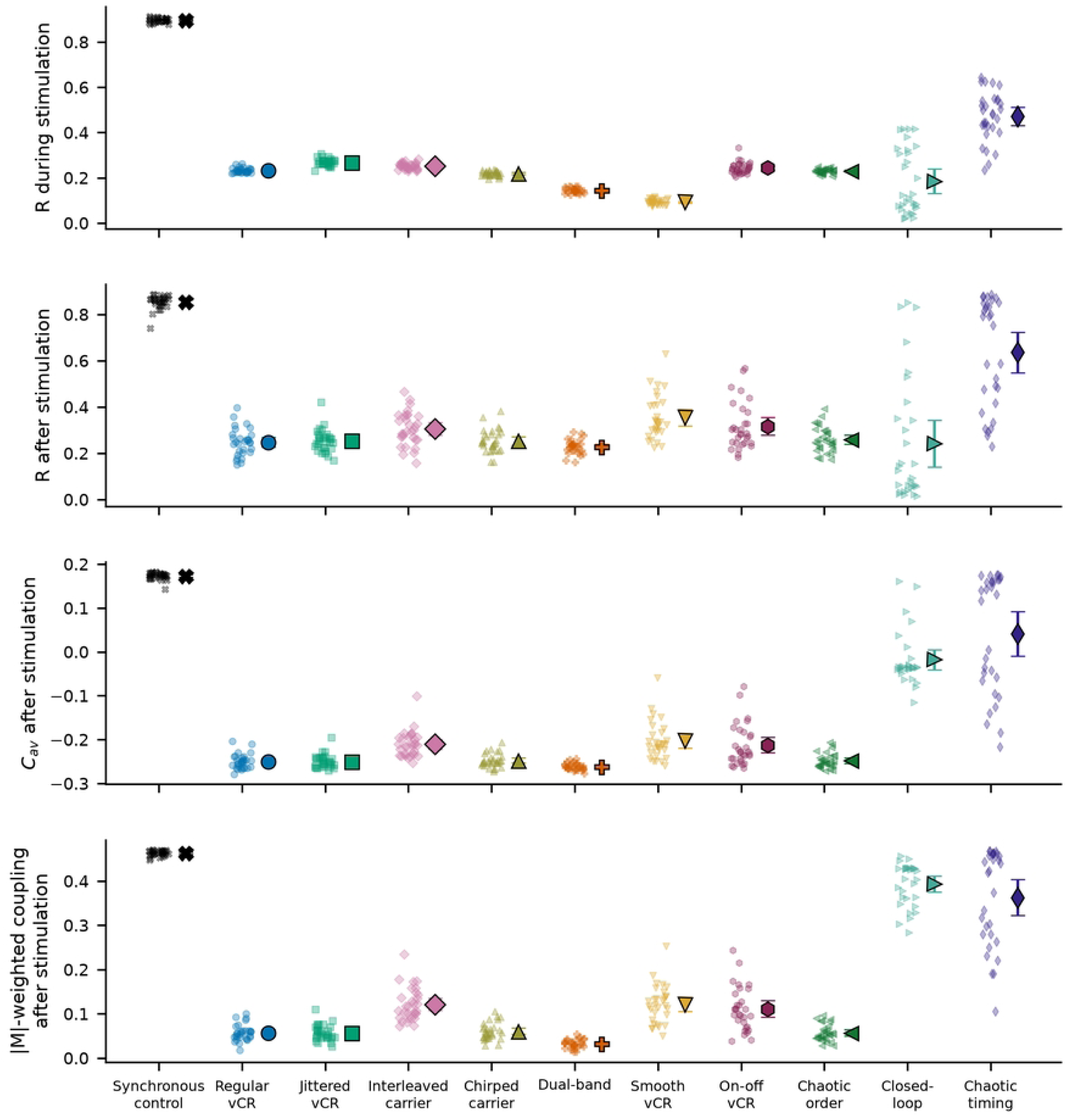
Outcome measures for all eleven patterns. *R*_on_, *R*_post_, *C*_av,post_ and |M|-weighted coupling after stimulation. Small symbols, individual networks (n = 30 per pattern); large symbols and bars, mean and 95% confidence interval.

**S2 Fig.**
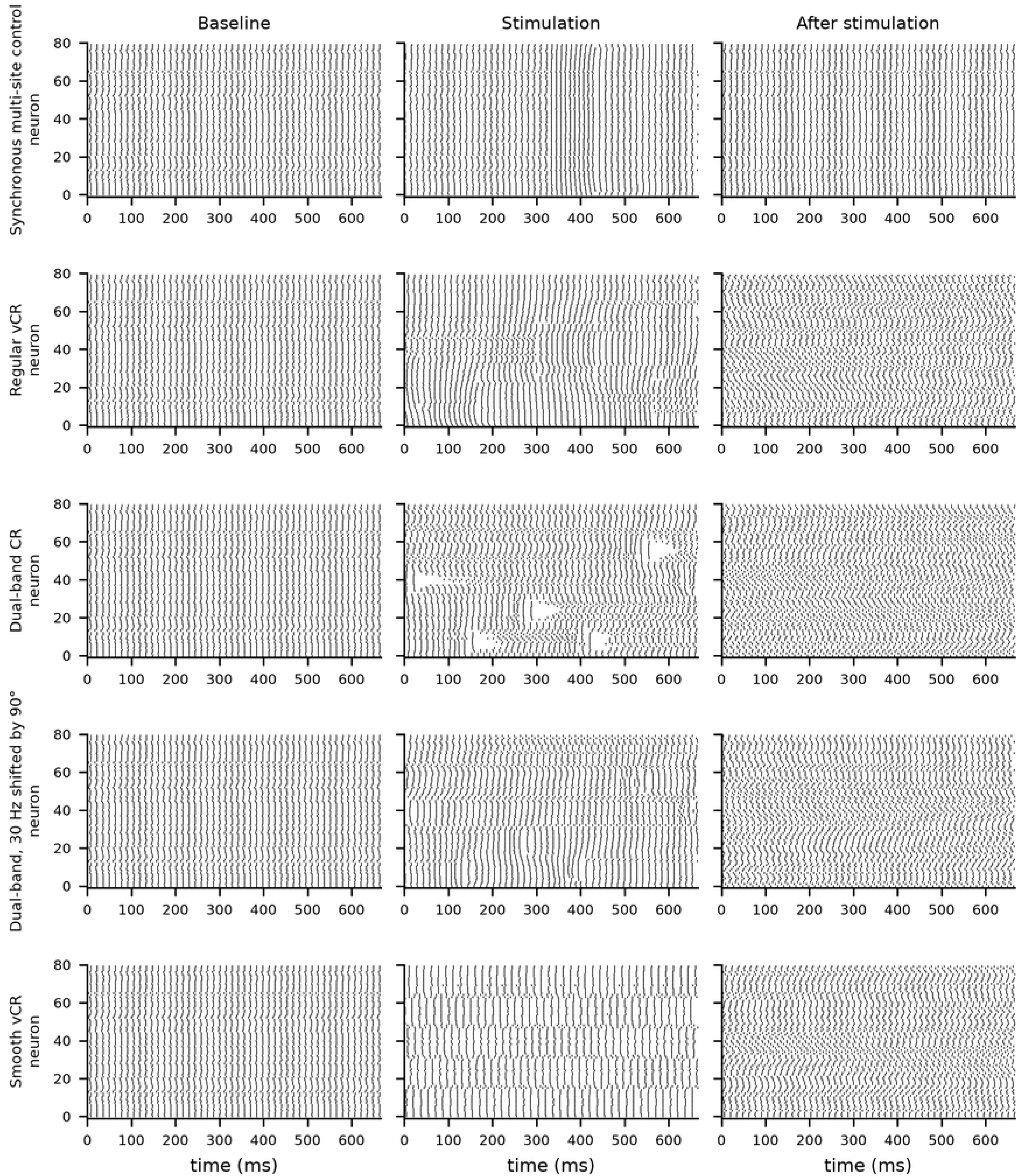
Staggered stimulation divides the synchronized network into site-specific clusters. Spike rasters of one network over one CR cycle (667 ms) at the end of baseline, 1 s after the onset of stimulation and 1 s after its offset, for the synchronous multi-site control, regular vCR, the dual-band burst, the dual-band burst with the 30 Hz component shifted by 90°, and smooth vCR.

**S3 Fig.**
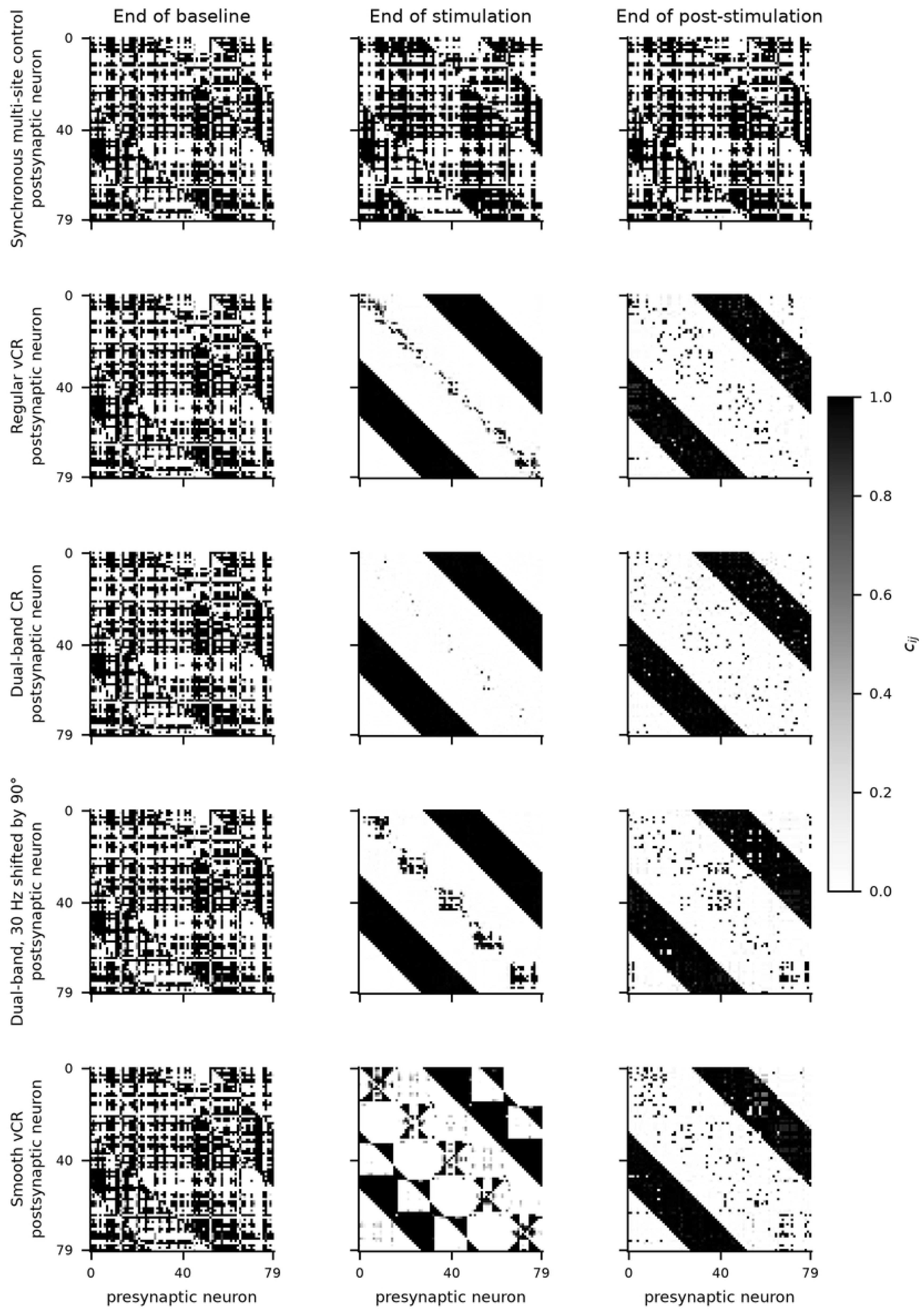
Staggered stimulation depresses local excitatory connections and potentiates distant inhibitory connections. Synaptic weight matrices *c*_ij_ of the same network at the end of baseline, at the end of stimulation and at the end of the post-stimulation period.

**S4 Fig.**
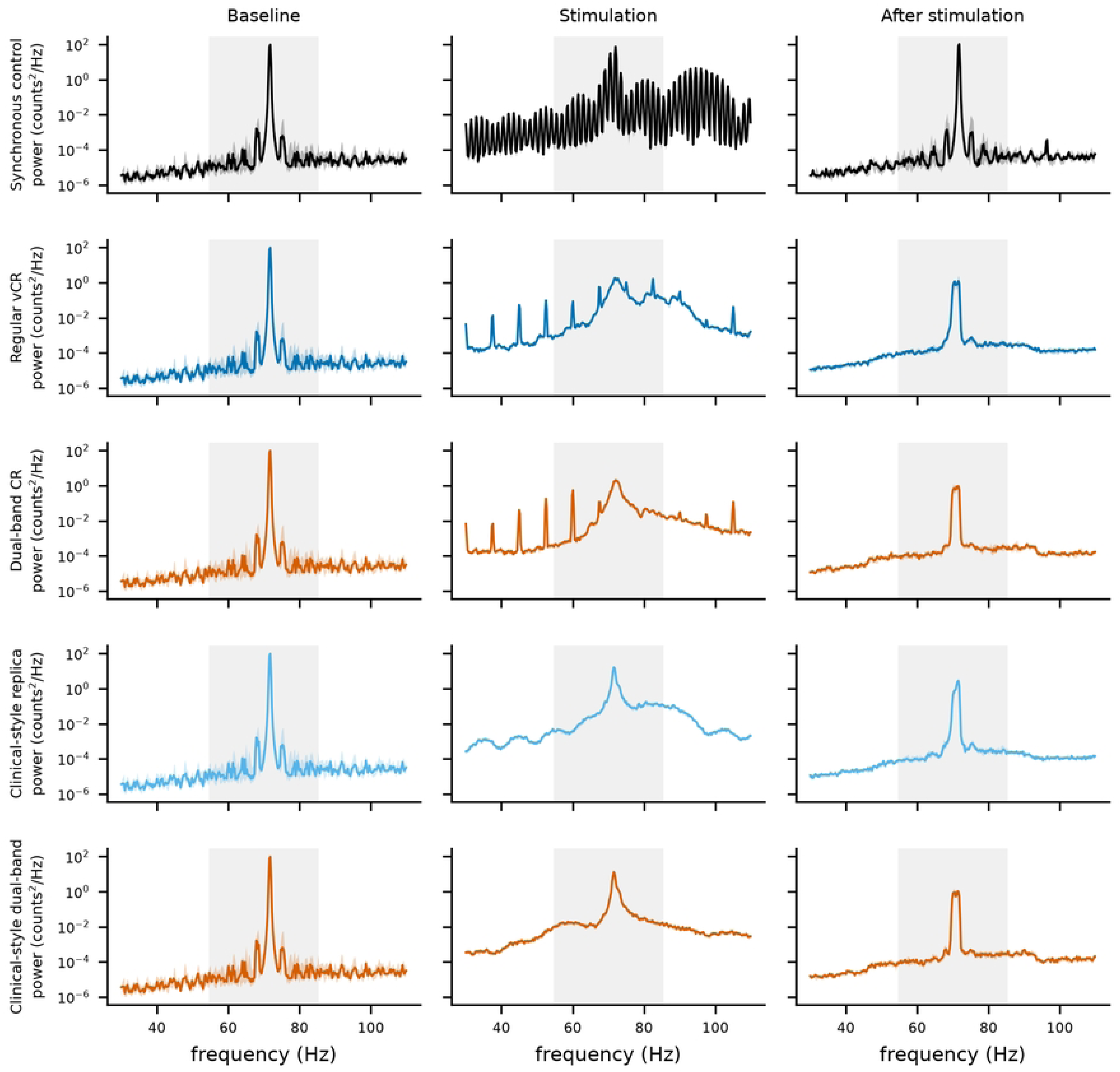
Population spectra during and after stimulation. Population spike-count spectra in 20 s windows at the end of baseline, at the start of stimulation and at the start of the post-stimulation period: median (line) and interquartile range (shading) over 15 networks; grey band, the 55 to 85 Hz synchrony band.

## Notes

### Competing Interest Statement

The authors have declared no competing interest.

